# Inosine Monophosphate Dehydrogenases are key players for functional development in Arabidopsis

**DOI:** 10.64898/2026.02.27.708476

**Authors:** Eva Dörfer, Zahra Sadeghi, Katharina Ebel, Adrian Heide, Vincent Frick, Tom Niehoff, Susanne Zehner, Lisa Fischer, Claus Peter Witte, Marco Herde, Torsten Möhlmann

## Abstract

In plants, nucleotide *de novo* synthesis is required throughout development to provide building blocks for the synthesis of DNA, RNA and nucleotide-derived cofactors. Inosine Monophosphate Dehydrogenases (IMPDH) are key enzymes in *de novo* guanylate synthesis. The genome of *Arabidopsis thaliana* encodes two IMPDHs and we identify IMPDH2 as the major isoform, being enzymatically active in the plant cytosol. Bimolecular fluorescence complementation (BIFC) analysis suggests that IMPDH1 and IMPDH2 interact with themselves and with Cytidine Triphosphate Synthetase (CTPS) isoforms, connecting purine and pyrimidine metabolism. Nucleotide quantification identifies IMPDH2 as bottleneck in guanylate biosynthesis, and plants lacking this isoform suffer from nucleotide imbalance and limitation, causing reductions in ribosomal RNAs by reducing Target of Rapamycin (TOR) activity. Impaired photosynthesis and growth were further consequences thereof. In contrast, strong overexpression of IMPDH1 supported growth and led to altered organ development in 10% of corresponding plants presumably due to altered auxin metabolism during embryo development.

**Significance statement:** This study reveals that Inosine Monophosphate Dehydrogenase 2 (IMPDH2) is required for early Arabidopsis development and growth. Knock-out mutants exhibit reduced amounts of metabolites (e.g. XMP, GMP, GTP) accompanied by reduced expression of genes in the GO-Terms Ribosome and Photosynthesis at seedling stage. This leads to reduced chlorophyll levels, impaired photosynthesis, and reduced growth. Marked overexpression of *IMPDH1* provoked abnormal organ development in 10% of cases presumably due to altered auxin metabolism.

## Introduction

Nucleotides are essential components of the genetic material and play important roles in cellular processes. They are involved in storage and transmission of genetic information in form of DNA and RNA. Further, nucleotides are needed as energy transmitters in many metabolic pathways and act as cofactors for sugar, starch, and phospholipid biosynthesis.

Plant purine *de novo* synthesis consists of 10 reaction steps in the plastid to generate inosine monophosphate (IMP). Two further plastidic reactions lead from IMP to AMP, while GMP is synthesized from IMP in the cytosol (Zrenner *et al*., 2006). Inosine Monophosphate Dehydrogenase (IMPDH) catalyzes the oxidation of inosine monophosphate (IMP) to xanthosine monophosphate (XMP) in the presence of NAD^+^. XMP is then aminated by Guanosine 5’-Monophosphate Synthetase (GMPS) using the amide group of glutamine as nitrogen donor and ATP for energetic coupling resulting in the production of GMP as well as glutamate, AMP, and pyrophosphate as side products (Figure 1). In a similar type of reaction, the final step in pyrimidine *de novo* synthesis is catalyzed by CTP-Synthetase (CTPS) performing the amination of uridine triphosphate (UTP) to cytidine triphosphate (CTP). There are five *CTPS* genes in Arabidopsis and the corresponding enzymes are all located in the cytosol (Moffatt and Ashihara, 2002, Zrenner *et al*., 2006, Witz *et al*., 2012, Daumann *et al*., 2018). The synthesis of CTP by CTPS relies heavily on GTP as allosteric activator which connects purine and pyrimidine biosynthesis (Levitzki and Koshland, 1972, Chang and Carman, 2008). Both IMPDH and CTPS can form filamentous structures named “cytoophidia” in vertebrates. These do interact with each other for regulation of protein activity and stability beyond the level of allosteric regulation (Chang *et al*., 2018).

**Figure 1.**
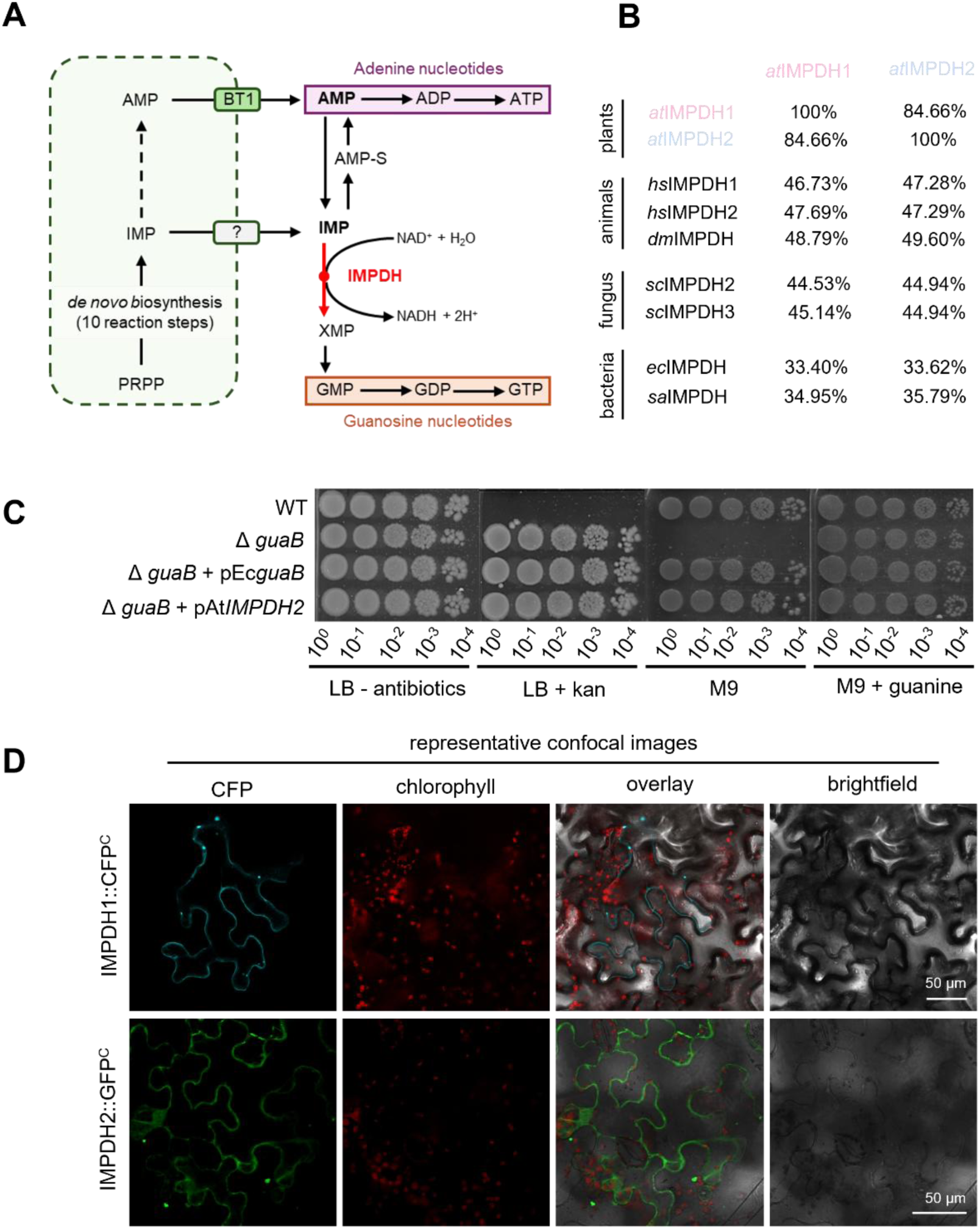
The Enzyme IMPDH is responsible for the synthesis of XMP. (**A**) Scheme of purine *de novo* synthesis. IMPDH catalyzes reaction of IMP to XMP under consumption of NAD+ and H_2_O. (**B**) Homology of Arabidopsis IMPDH to corresponding proteins from selected animals, fungi and bacteria. Percentage of similar amino acid residues is given, based on protein alignments. at, *Arabidopsis thaliana*; hs, *Homo sapiens*; dm, *Drosophila melanogaster*; sc, *Saccharomyces cerevisae*; ec, *Escherichia coli*; sa, *Staphylococcus aureus*. (**C**) Spot test for AtIMPDH2 activity in complemented Δ*guaB E. coli* strains. (**D**) Localization of *IMPDH1::CFP* and *IMPDH2::GFP* transiently expressed in *N. benthamiana*.

Human IMPDH exists as two isoforms (Carr *et al*., 1993) and mutations in both can lead to diseases. Therefore, vital interest in IMPDH biology results from the observation of cell proliferation and tumor growth in humans correlating with increased purine *de novo* synthesis and specifically IMPDH activity, making IMPDH a target for cancer therapy (Jackson *et al*., 1975). The retina is characterized by a high demand for purine nucleotides and mutations in human IMPDH1 cause retinal degeneration (Bowne *et al*., 2002). Mutations in human IMPDH2, however, have been associated with a neurodevelopmental disease (Burrell and Kollman, 2022). Thus, both human isoforms exhibit unique but different physiological functions.

Although IMPDH also exists in two isoforms in many animals and plants, Drosophila (*Drosophila melanogaster*) only harbors one isoform (Sifri *et al*., 1994) and the same holds true for plants such as Rice (*Oryza sativa*), Barley (*Hordeum vulgare*), Potato (*Solanum tuberosum*) and Wine (*Vitis vinifera*), to name a few (TAIR, The Arabidopsis Information Resource, www.arabidopsis.org). Apparently, a physiological diversification of IMPDH is not required *per se*.

In its monomeric form, IMPDH has two domains, a catalytic domain which binds the substrates IMP and NAD^+^, and a regulatory domain which can bind adenine and guanine nucleotides at distinct sites, thereby altering the oligomeric state and activity of IMPDH (Burrell and Kollman, 2022). Assembly of IMPDH cytoophidia reflects an upregulation of IMPDH activity correlating with rapid cell proliferation. From this it was concluded that maintenance of the GTP pools requires IMPDH filamentation (Keppeke *et al*., 2018).

In plants, purine *de novo* synthesis proceeds in plastids and results in the export of AMP by the BT1 transporter. It has been speculated that inosine monophosphate (IMP) can be directly transported into the cytosol by a yet unknown transporter (Leroch *et al*., 2005, Witte and Herde, 2020) since the IMPDH and GMPS reactions for GMP synthesis are located in the cytosol. IMPDH from plant tissue has first been described and purified from soybean (Cao and Schubert, 2001) and two putative IMPDH isoforms have been identified by homology in Arabidopsis, IMPDH1 (At1g79470) and IMPDH2 (At1g16350), both sharing 84% identity at amino acid level with another (Collart *et al*., 1996, Witte and Herde, 2020, Maekawa *et al*., 2024).

Here, we describe knock-out and overexpression mutants for both *IMPDH* isoforms and reveal a major role of IMPDH2 during ribosome biogenesis, and the establishment of photosynthesis while IMPDH1 is required for functional embryo development and cotyledon formation. Furthermore, we present subcellular localization studies by fluorescent fusion proteins and describe interactions by bimolecular fluorescence complementation (BIFC) with Cytidine Triphosphate Synthase (CTPS) enzymes. Transcriptomic analysis reveals alterations in gene expression in four- and eight-day-old *IMPDH2* knock-out lines, complemented by nucleotide metabolite quantification in seedlings that were obtained from imbibed seeds transferred to growth conditions for 12h, 48h or for 8d.

## Results

### Phylogenetic relations among IMPDH family members

The central position of IMPDH in purine metabolism suggests a vital role of this enzyme in the establishment of purine nucleotide homeostasis (Figure 1A). The high similarity between both Arabidopsis isoforms of 85% suggests a redundant function in XMP synthesis. Similarities to fungal and animal homologs are in the range between 44.5% to 49.6% and thus still high, whereas the similarity of *At*IMPDH to bacterial IMPDH ranges between 33.40% to 35.79% (Figure 1B). Phylogenetic analysis reveals clustering of IMPDH isoforms within Brassicaceae, however in all other plant species analyzed, the isoforms 1 and 2 are more closely related to each other compared to any IMPDH from another species (Figure S1) also supporting the idea, that Arabidopsis has two IMPDH isoforms because they have different, for example tissue specific functionalities. Some plants like *Manihot esculenta*, *Nicotiana tabacum*, *Oryza sativa* only harbor a single IMPDH. In animals with mostly two isoforms, IMPDH1 and IMPDH2 form distinct clusters (Figure S1). Furthermore, algal and fungal IMPDHs are more closely related to the plant cluster than bacterial IMPDHs. (Figure S1).

### IMDPH2 is enzymatically active

Enzymatic activity of IMPDH2 was assessed in an *Escherichia coli* (*E. coli*) complementation approach using a mutant of the *E. coli* IMPDH encoded by *guaB*. A Δ*guaB* knock-out strain (JW5401-KC, Keio Collection) is unable to grow on minimal medium because GMP *de novo* synthesis is compromised. This mutant was complemented with either *guaB* from *E. coli* (Δ*guaB* + pEc*guaB*), or with the codon optimized *IMPDH2* gene from *A. thaliana* (Δ*guaB +* pAtIMPDH2opt). For genotyping of the corresponding *E. coli* strains see Figure S2. All tested strains grew on control M9 minimal medium with 0.5 mM guanine which can be salvaged to GMP. However, on M9 medium without guanine the Δ*guaB* strain failed to grow. This defect could be complemented by *guaB* (strain Δ*guaB* + pEc*guaB*) and importantly also by *IMPDH2* (strain Δ*guaB +* pAtIMPDH2opt.) revealing that AtIMPDH2 is enzymatically functional (Figure 1C).

### Both IMPDH isoforms locate to the cytosol and can interact with themselves and with Cytidine triphosphate Synthase (CTPS)

AMP and presumably also IMP are exported from plastids for the further synthesis of adenylates and guanylates (Leroch *et al*., 2005, Zrenner *et al*., 2006, Witte and Herde, 2020). As IMPDH isoforms do not carry N-terminal extensions that could serve as organellar targeting sequence, it is likely that they are located in the cytosol. To test this assumption, IMPDH1 and IMPDH2 were synthesized as fusion proteins with C-terminal Cyan Fluorescent Protein (CFP) and Green Fluorescent Protein (GFP), respectively, using transient expression in *Nicotiana benthamiana*. CFP and GFP signals were observed at the cell margin in epidermal leaf cells which is indicative of a cytosolic localization because the cytosol surrounds the extensive central vacuole in these cells. Often, fluorescence appeared in macromolecular structures, which we denominate polymers from now on (Figure 1D, Figure S3).

IMPDH and CTPS catalyze key reactions in the synthesis of the pyrimidine ribonucleotides GTP and CTP. For this, CTPS enzymes from animals and plants require allosteric activation by GTP (Lunn *et al*., 2008, Daumann *et al*., 2018), which functionally links purine and pyrimidine *de novo* synthesis. Furthermore, protein-protein interactions between CTPS and IMPDH have been identified as additional regulatory mechanisms in human cells where these enzymes were found in cytoophidia as rods and rings (Carcamo *et al*., 2011, Keppeke *et al*., 2018). As indications for cytoophidia formation of plant CTPS isoforms were reported as well (Daumann *et al*., 2018, Alamdari *et al*., 2021, Krämer *et al*., 2022), we were curious about interactions between IMPDH isoforms themselves and with CTPS variants. To test this hypothesis, YFP N-and C-terminally halves were translationally fused to IMPDH and CTPS N-and C-termini for bimolecular fluorescence complementation (BiFC) analysis. We observed that IMPDH1 interacts with IMPDH2 and all four tested CTPS isoforms. IMPDH2 on the other hand only interacts with CTPS1-3 (Figure 2). In case of IMPDH1 only the interaction with CTPS3 results in formation of filaments, whereas IMPDH2 interaction with CTPS1, 2 and 3 promoted filament formation (Figure S4).

**Figure 2.**
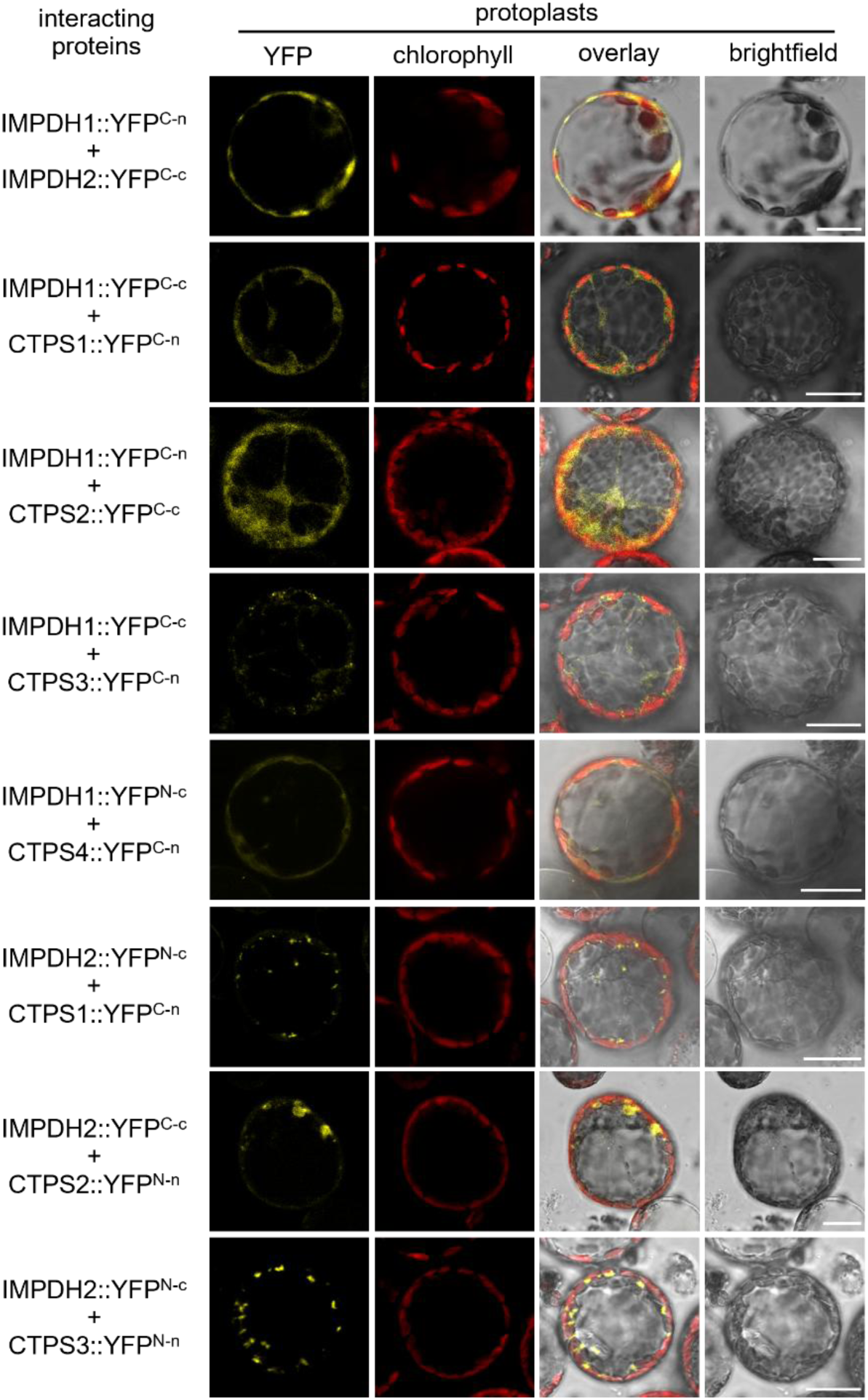
Bimolecular Fluorescence complementation (BIFC) of IMPDH isoforms and CTPS isoforms. Shown are interaction of combinations of each two different isoforms after transient expression in *N. benthamiana* leaves. IMPDH and CTPS proteins were fused C- or N-terminally with either a C- or N-terminal YFP fragment, as indicated. Confocal laser scanning microscopy was applied for imaging. Scale bar = 20 µm

To obtain clues about the functional state of the observed polymers, *IMPDH2* was transiently expressed in tobacco leaves. Transfected leaf discs were incubated with various metabolites. These were the substrate of IMPDH, IMP, the IMPDH-specific inhibitor mycophenolic acid (MPA), ATP, which produces an active conformation of IMPDH in humans, and GTP, which triggers feedback inhibition in humans (Buey *et al*., 2015, Johnson and Kollman, 2020), and GTP together with ATP. Under control conditions, as observed before, only a few polymers were monitored. Neither IMP, MPA nor GTP addition triggered polymer formation. Interestingly, when ATP was added, many polymers could be observed, and the enzyme was no longer present in dissolved form. When GTP and ATP were supplemented simultaneously, the ATP-induced polymer formation was noticeable, with IMPDH2 also partially returning to a dissolved state (Figure S5).

### IMPDH2 mutants show severe developmental defects

To obtain insights into the physiological function of Arabidopsis IMPDH isoforms, T-DNA mutants for both were identified. These are the homozygous mutants of *IMPDH1* (SAIL_5_B10C; *impdh1-1*) and *IMPDH2* (SALK_008653; *impdh2-0* and SALK_201269; *impdh2-1*). Corresponding transcript levels of the mutated *IMPDH* gene were not detected, while transcripts of the respective other isoform were reduced as well to about 50% wild-type level (Figure S6 A-C). Moreover, complementation lines for *impdh1-1* and overexpression lines for both IMPDH isoforms were generated. Three *IMPDH1* complementing lines (*IMPDH1* c#1-3) with transcript levels close to WT were identified (Figure S6D). Furthermore, three *IMPDH1* overexpression lines (*IMPDH1* ox#1-3) with medium to high overexpression (10-, 21-, and 46-fold relative to WT) were generated. For *IMPDH2,* three lines (*IMPDH2* ox#1-3) with medium to high overexpression (4-, 14-, and 33-fold relative to WT) were obtained (Figure S6E,F).

All mutant plant lines analyzed in this work were able to complete a full lifecycle. Due to the high similarity of both IMPDH proteins we speculated about a functional redundancy. Crosses between *impdh1-1* and *impdh2-0* resulted in 10.5 % wildtypes, 31.6 % heterozygous (het) plants for both alleles and 42.1 % homozygous (hz) *IMPDH2* mutants (Supplemental Table S1). No homozygous double knock-out plants could be observed, indicating that the homozygous loss of both *IMPDH* isoforms is embryo lethal, as previously described (Maekawa et al., 2024). In line with this, aborted seeds in developing siliques were already observed in both single *IMPDH* knock-out mutants at rates of 27,3% for *impdh1-1* and 25,4% for *impdh2-0.* Seed abortion was accompanied by reduced length of siliques from both knock-out lines (Figure S6G-H).

In *impdh1-1*, and in both *impdh2* mutant lines, reduced FW levels were seen, accompanied by severe chlorosis in both *impdh2* lines after 4 days of growth on soil (Figure 3A). While plant size was still reduced after 8 d of growth, the leaves of *impdh2* lines were not chlorotic anymore. After 21 d, *impdh1-1*, *impdh2-0* and *impdh2-1* still showed a reduced rosette size (Figure 3A). The differences in plant size throughout growth were also resembled in the corresponding fresh weights (FW). During 4 d to 21 d of growth, all *impdh* ko lines had a lower FW compared to WT. While 4-day-old *impdh1-1* showed 14% lower FW compared to WT, in *impdh2-0* and *2-1* the reduction was 22% and 19%, respectively. After 8 days of growth FW reductions were 11% for *impdh1-1*, and app. 40% for both *IMPDH2* mutant lines (Figure 3B). Opposite to this, the IMPDH1ox#2 mutants where larger after 8 days of growth (Figure S7 A) and had a slightly higher FW (8%, Figure 3B). The chlorotic *impdh2-0* and *impdh2-1* contain only approximately one third of chlorophyll compared to the WT after 4d of growth. These deficits adjust to the WT chlorophyll level after 8 d (Figure 3C).

**Figure 3.**
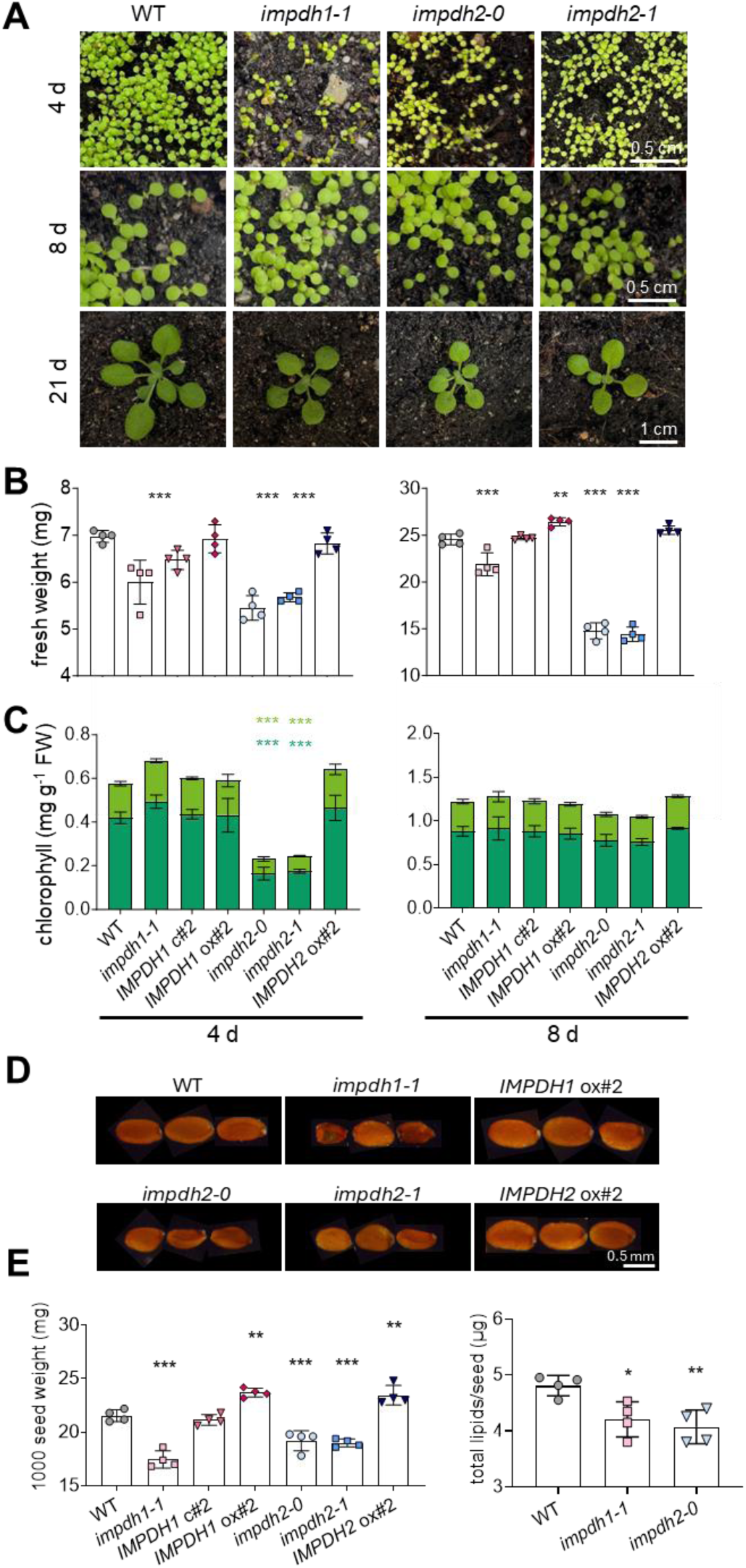
Phenotypical analysis of *IMPDH* loss-of-function mutants. (**A**) Phenotypic analysis of WT, *impdh1-1*, *impdh2-0* and *impdh2-1* Arabidopsis plants. Plants were grown under standard conditions (21 °C day and night temperature, 10 h day length and 120 *µ*E light intensity) for different length of time as indicated. (**B**) Fresh weights of WT, *impdh1-1*, *IMPDH1* c#2, *IMPDH1* ox#2, *impdh2-0*, *impdh2-1* and *IMPDH2 ox#2* after 4 d and 8d of growth. (**C**) Chlorophyll a and b levels of the same plants as in (B) after 4 d and 8d of growth. (**D**) Typical appearance of mature seeds from *IMPDH* mutants. (**E**) 1000 seed weight and total lipids per seed of WT and *IMPDH* mutants. Error bars represent mean values ± SD. Asterisks indicate significant differences between WT and mutant using a one-way ANOVA followed by a dunnett’s multiple comparisons test: ** p-value ≤ 0.01, ***p-value ≤ 0.001

Mature seeds of both *IMPDH* knock-out mutant lines were smaller and showed deformations compared to WT seeds. Furthermore, the seed weight of *impdh1-1* and *impdh2-0* was lower compared to WT seeds, whereas the corresponding overexpression seeds were heavier and larger (Figure 3D,E). As Arabidopsis seeds contain up to 46% lipids (Holzl and Dormann, 2019), the total lipid content was quantified. We observed reduced total lipid contents based on seed weight for *impdh1-1* and *impdh2-0* (Figure 3F).

As chlorosis was observed in *IMPDH* mutants which is an indicator for impaired photosynthetic effectiveness, we performed pulse amplitude modulation (PAM) measurements. Here, we could detect a massive decrease of PS yield (Fv/Fm) at 4 d of growth for *impdh2-0* and *impdh2-1*. After 8 d Fv/Fm was still slightly reduced in these seedlings. After 21 days of growth, Fv/Fm reduction was restricted to the new developed leaves in both *impdh2* mutants, indicating a vital function of IMPDH2 during cell proliferation as proliferating tissues have a high demand for nucleotide supply (Fig 4A,B). Additionally, transcript levels of photosynthesis associated genes were reduced in 4-day-old *impdh2-0* seedlings (Figure 4C). Especially *psaB* was massively reduced in transcript amount. When inspecting protein contents of IMPDH mutants it appeared that in *impdh2-0* seedlings a lower extractable amount of protein was apparent and reduced RUBISCO levels largely contributed to this (Figure 4D). Steady state levels of photosynthesis proteins (PsaB, PsbA, and LHCB2) were reduced as well, in line with the observed reductions in corresponding transcript levels (Figure 4D).

**Figure 4.**
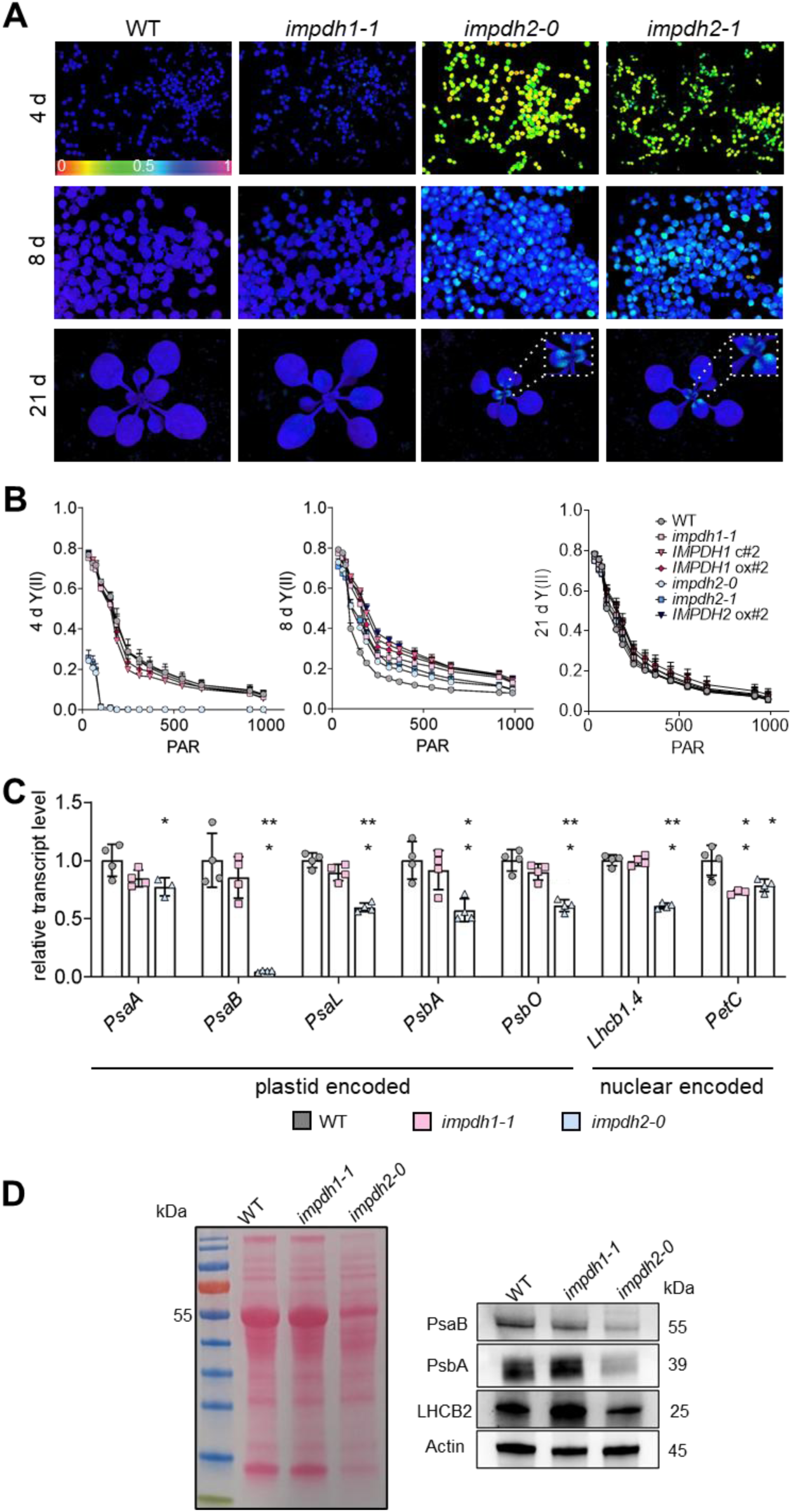
Photosynthetic alterations in *impdh2* mutants. (**A**) Representative PAM images of WT, *impdh1*, *impdh2-0* and *impdh2-1* are depicted for maximum PSII capacity (Fv/Fm). (**B**) Light curves of Y(II) for WT and *IMPDH* mutants after 4, 8 and 21 days of growth. (**C**) Expression levels of photosynthetic genes in *impdh1-1* and *impdh2-0* after 4d of growth. (**D**) Ponceau staining of total extracted protein corresponding to 4 mg FW per genotype (4-d-old seedlings). Immunoblot of photosynthesis associated proteins in 4 d old seedlings with Actin as loading control and corresponding ponceau staining. Error bars represent ± SD. Asterisks indicate significant differences between WT and mutant using one-way ANOVA followed by dunnett’s multiple comparisons test: * p-value ≤ 0.05, ** p-value ≤ 0.01, ***p-value ≤ 0.001

### Morphological alterations in *IMPDH1* over expressors may result from altered auxin metabolism

In the strong overexpression lines *IMPDH1 ox#2* and ox#3 besides slight growth improvements morphological alterations were observed. About 10% of *IMPDH1* ox#2 and *IMPDH1* ox#3 plants showed abnormal leaf numbers. At seedling stage apparently 3 or 4 cotyledons were observed, and these plants then also grew 3 or 4 new leaves at a time (Figure 5A). Growth of one petiole that divides into leaves was seen in these plants as well. Similar effects were seen for siliques, where two siliques grow from the same pedicel (Figure 5A). Additional cotyledons and -alterations were also observed in mutants for PIN1 and PINOID proteins involved in auxin metabolism (Furutani, 2004) Therefore, we speculated that imbalanced auxin metabolism during embryo development might account for the observed improper organ development. Levels of auxin responsive genes in embryos 6 DAF were quantified, and it appeared that transcripts of *SAUR9* and *SAUR10* (small auxin responsive 9 and 10) were markedly upregulated in *IMPDH1* ox#2 (Figure 5C). This increase in *SAUR* transcripts indicates massive overproduction of auxin in the *IMPDH1 ox#2* embryos. *GH3.3*, on the other hand, is an auxin conjugation enzyme that breaks down excess auxin. The increased expression of *GH3.3* confirms the accumulation of auxin in the *IMPDH1 ox#2* embryos, whereas the reduced transcript content of *GH3.3* in *impdh1-1* embryos indicates a reduced amount of auxin. The expressions of the transcription factor ‘Auxin-Responsive Factor 5’ (ARF5) and the auxin repressor IAA12 also show typical responses to increased auxin levels in the *IMPDH1 ox#2* embryos. ARF5 activates auxin signaling and developmental genes. *IAA12* binds to *ARF5*, thereby blocking it (Hamann, 2002). However, increased auxin levels lead to the repression of *IAA12*, so the regulation of auxin-responsive genes by *ARF5* can take place again (Maraschin Fdos *et al*., 2009). The transcript levels of the genes *PIN1*, *Wuschel-Related-Homeobox 2* (*WOX2*) and *WOX8* were examined to obtain information on the distribution of the phytohormone. The reduction in *PIN1* expression in the *IMPDH1* ox#2 embryos indicates reduced auxin transport in the basal part of the embryo. This is consistent with the increase of *WOX2* expression, which is in the apical part of the embryo, and decreased *WOX8* expression in the *IMPDH1* ox#2 embryos, which* is needed for cell fate determination in the basal part of the embryo. Taken together these auxin imbalances in the embryo likely contribute to altered organ development.

**Figure 5:**
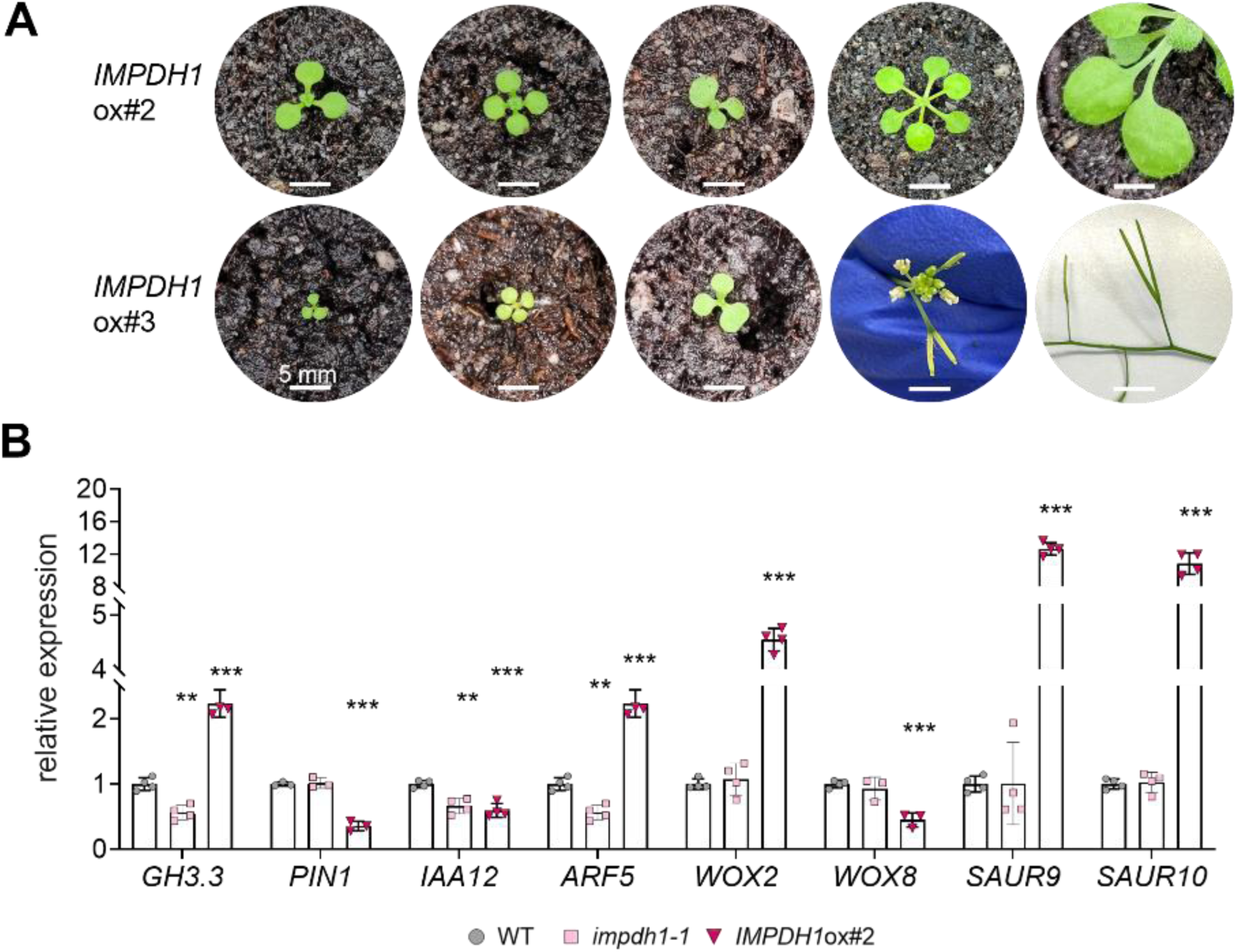
Phenotypical analysis of *IMPDH1* overexpression mutants. (**A**) Phenotypic analysis of *IMPDH1* ox#2 and ox#3 Arabidopsis plants showing alterations in cotyledon number and shape as well as altered silique growth. (**B**) Gene expression levels of auxin related genes in WT, *impdh1-1* and *IMPDH1* ox#2 embryos harvested 6 days after fertilization (DAF). Error bars represent ± SD. Asterisks indicate significant differences between WT and mutant using a one-way ANOVA followed by a dunnett’s multiple comparisons test: * p-value ≤ 0.05, ** p-value ≤ 0.01, ***p-value ≤ 0.001

### Nucleotide metabolism is altered in *IMPDH* mutants

As we speculated about limiting guanylate supply as reason for the low chlorophyll content and the reduced FW in *IMPDH2* knock-out mutants, a rescue experiment was performed. For this, WT, *impdh2-0* and *IMPDH2* ox#2 seedlings were grown in liquid culture with GMP, guanine or mycophenolic acid (MPA), an IMPDH specific inhibitor. With GMP supplementation, *impdh2-0* seedlings lose their chlorotic phenotype and chlorophyll levels resemble those of the WT whereas the supplementation with guanine leads to an increase in FW in *impdh2-0* to WT level under standard conditions (Figure S8). MPA dependent reduction of chlorophyll levels was less pronounced in *IMPDH2* ox#2 (Figure S8), supporting our initial hypothesis. Histological stainings of *proIMPDH2* ::GUS lines revealed staining of cotyledons, shoot apical meristem and roots in six day old seedlings. When incubated with guanine or GMP, staining of cotyledons appeared slightly reduced. In contrast, when the IMPDH inhibitor MPA was applied, *proIMPDH2::GUS* staining clearly increased. Thus, *IMPDH2* expression seems to respond inversely with respect to guanylate availability (Figure S8).

IMPDH is central for the guanylate biosynthesis pathway involving the nucleotides IMP, XMP and GMP. Plants with modified IMPDH activity might therefore be affected in the content of these nucleotides, which may cause further alterations of general nucleotide homeostasis. To test this hypothesis, we quantified nucleotides in germinating seeds and seedlings by LC-MS analysis at several developmental time points: at the end of imbibition period at 4°C and after 12 h, 48 h and 8 days of growth. Directly after imbibition neither IMP nor XMP were detectable but the content GMP was reduced strongly in *impdh2* seeds and also slightly in *impdh1-1* seeds, whereas this was not the case in the overexpression lines (Figure 6). In principle, the same was observed for GTP but with higher statistical uncertainty (p > 0.05). At the later time points, IMP and XMP became detectable in seedlings. The IMP content was generally higher and the XMP content lower in *impdh2* seedlings while this was not the case in the *impdh1-1* line. Similarly, GMP and GTP concentrations were only reduced in the *impdh2-0* line. The *IMPDH2* ox#2 complemented the metabolic phenotype of the corresponding mutant at all time points, sometimes even over-compensating the mutant by reducing the IMP content below wild type level (at 12 hours) and increasing the XMP concentration above that of the wild type (at 8 d and in tendency, but with p > 0.05, also at 48 h). The metabolite content of *IMPDH1* ox#2 and the wild type did not differ at any time point. Interestingly, the metabolic pattern for GMP and GTP in the different *IMPDH* variants was in principle mirrored by the other nucleotides (AMP and ATP, UMP and UTP, CMP and CTP; (Figure S9). The statistical support for differences in individual metabolites is often weak, but overall, the nucleotides in the different *IMPDH* variants show similar trends.

**Figure 6:**
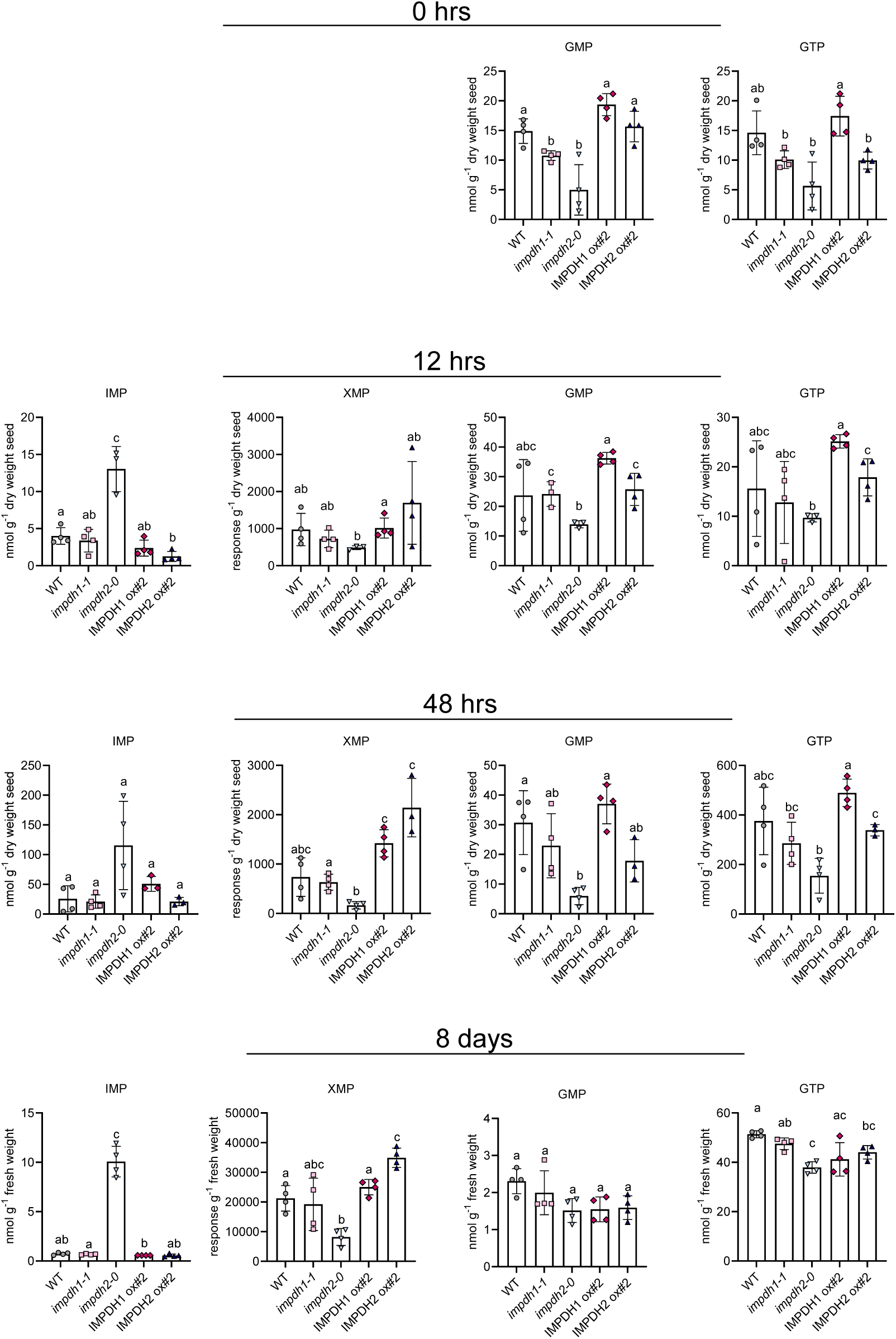
Metabolic analysis of guanylate biosynthesis pathway nucleotides in genetic *IMPDH* variants over a developmental time course. Nucleotides quantified by LC-MS after imbibition (0 h) and after 12 h, 48 h and 8 d of growth. Three or four biological replicates were analyzed, error bars are SD. The statistical analysis used the two-sided Tukey’s multiple pairwise comparison test with the sandwich variance estimator. Different letters indicate p values < 0.05.

The metabolite data indicate that IMPDH2 is the more important enzyme overall, which agrees with its generally higher expression compared to IMPDH1. However, in the imbibed seed, IMPDH1 clearly contributes to the overall IMP to XMP turnover. Changes in the guanylate content by alteration of IMPDH activity results in parallel changes of other nucleotides presumably by regulation of nucleotide homeostasis that has also been observed in other cases (Straube *et al*., 2021)

### Transcriptome analysis of chlorotic *impdh2-0* seedlings reveals changes in gene expression in the categories “ribosomal proteins”, “nucleotide metabolism” and “photosynthesis”

To reveal the consequences of nucleotide limitation on changes in gene expression in *impdh2* mutants, an RNASeq analysis was performed. In this analysis, *impdh2-0* seedlings were compared to WT controls of the same size early after germination. In one comparison, 4-day-old *impdh2-0* seedlings (showing chlorosis) were compared to 3-day-old WT seedlings. Furthermore, slightly older seedlings (*impdh2-0* 8 day-old and WT 6-day-old) were compared. Here the *impdh2-0* seedlings had already overcome chlorosis. All samples were harvested and analyzed in three independent biological replicates.

In the comparison *impdh2-0 4d* vs WT 3d, 3.243 differently expressed Genes (DEGs; adjusted p_adj._-value<0.05) were identified. Among these 1.472 were upregulated and 1.771 downregulated (Figure 7A). When comparing *impdh2-0 8d* vs WT 6 d, only 985 DEGs were identified, app. half of them were up- and the other half downregulated (Figure 7B). Between 8d vs 4d post germination of *impdh2-0,* 4917 DEGs occurred, 2491 down and 2426 upregulated (Figure 7C). GO-term analysis revealed that ribosome, translation, photosynthesis and carbohydrate metabolic process are the most significantly altered GO-Terms in 4d old *impdh2-0* seedlings. When comparing 8d old *impdh2-0* seedlings to WT of a comparable developmental stage, all three named categories do not appear as significant GO-terms anymore (Figure 7D,E). Within the category “ribosome” 157 altered ribosomal genes were identified in our analysis, of these 17 genes were up- and 140 were down regulated (Figure 7F). Among the upregulated ribosomal proteins 11 were chloroplast located, including AT3G22450 (log2fold change 2.78) which encodes ribosomal protein uL18-L1. This protein is required for proper splicing of mitochondrial introns and loss of function mutants show a dwarf phenotype (Wang *et al*., 2020). One of the downregulated transcripts encodes for Ribosomal Protein S6A (RPS6A), which is phosphorylated by S6K. Prior, S6K is phosphorylated by TOR (Kim *et al*., 2014). RPS6A is required for proper translation and may affect in addition histone deacetylation (Kim *et al*., 2014).

**Figure 7:**
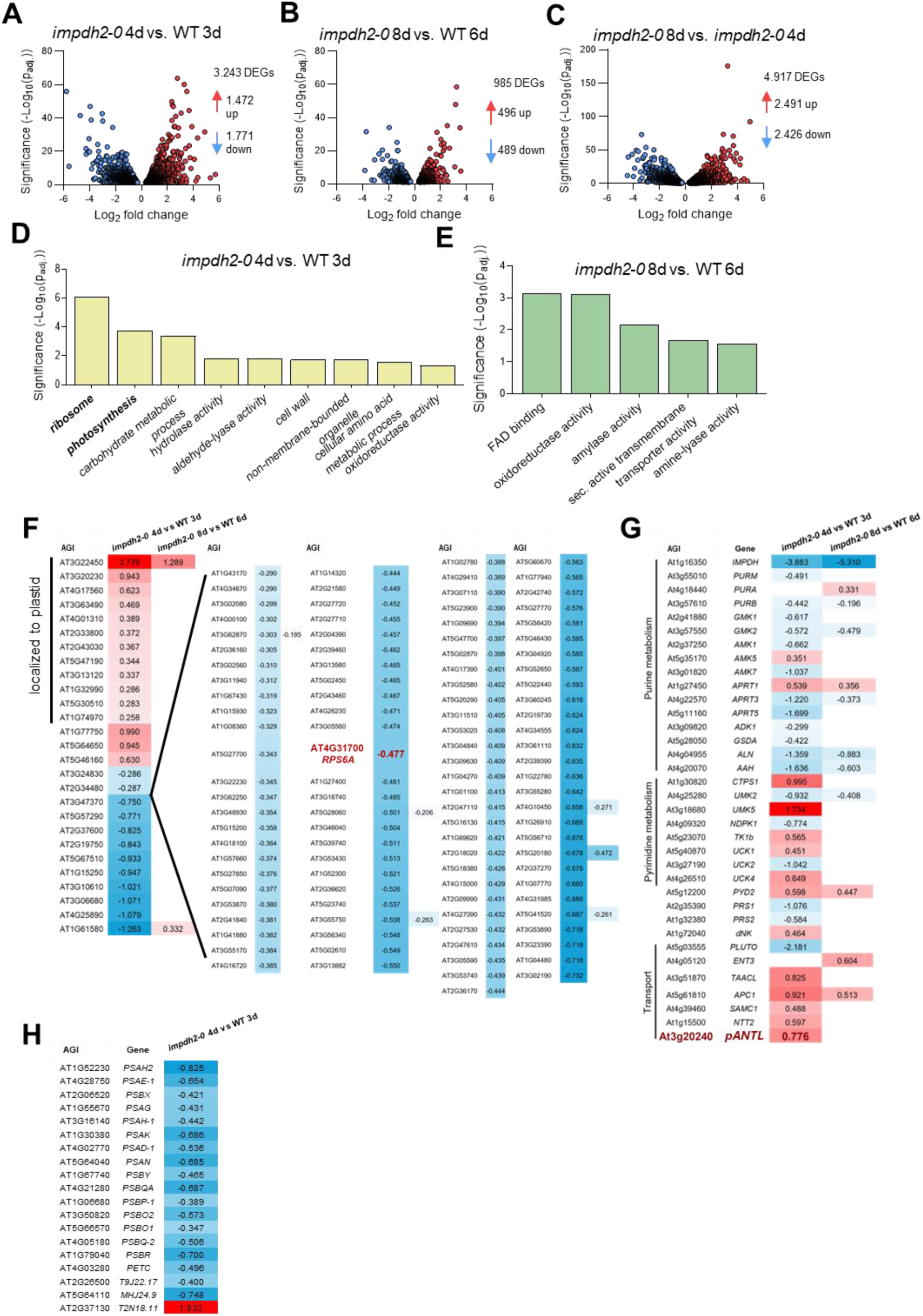
Transcriptome analysis of *impdh2-0* and corresponding WT seedlings at two developmental stages. (**A-C**) Volcano plots of DEGs (*impdh2-0* vs WT) and seedlings at same developmental stages, during their early chlorotic phase (4 d for *impdh2-0*) and the compensated phase (8 d for *impdh2-0*). (**D**-**E**) GO-term analysis of transcriptomic differences measured between *impdh2-0* and WT (**F**-**H**) Heat map analysis of transcriptomic differences measured between *impdh2-0* and WT seedlings of DEGs from the GO-terms: Ribosome, Photosynthesis and Nucleotide metabolism. Numbers indicate log_2_-fold differences.

When inspecting “nucleotide metabolism” related genes in the comparison *impdh2-0* 4d vs WT 3d, most DEGs in purine metabolism were down, with the exception of those encoding chloroplast localized proteins APRT1 and AMK5 catalyzing salvage of purine nucleobases (Witte and Herde, 2020) (Figure 7G). In pyrimidine metabolism, an opposite trend was observed as many genes in this category were upregulated. Uridine monophosphate kinase 5 (UMK5) represents the most strongly upregulated gene (1.7 log2 fold). Besides acting as kinase, UMK5 (PUMPKIN) can serve as RNA binding protein affecting plastid translation and splicing (Schmid *et al*., 2019). Interestingly, also several organellar transporters were upregulated as well, possibly to support nucleotide and energy balancing between compartments (Figure 7G). Eight-day old *impdh2-0* plants clearly showed a more WT like gene expression (Figure 7G). In the category Photosynthesis, all identified DEGs showed downregulation in *impdh2-0 4d* vs WT 3d, comprising genes which encode subunits of PSI and PSII and expression of all these increased again in the comparison *impdh2-0* 8d vs WT 6d (Figure 7H).

## Discussion

In this work we could show that Arabidopsis harbors two isoforms of inosine monophosphate dehydrogenase (IMPDH) and both locate to the plant cell cytosol. Moreover, in a complementation approach, IMPDH2 was shown to actively convert IMP to XMP, a key step in guanylate biosynthesis (Figure 1), and most likely this is also true for IMPDH1, based on the phylogenetic analysis (Figure S1). However, the most pronounced phenotypic alterations in this study were observed in *IMPDH2* knock-out mutants, which showed a clear developmental delay 4 days after germination. As *IMPDH2* shows much higher expression levels compared to *IMPDH1* in almost all tissues, this is indicative of an important function of the conversion of IMP to XMP in early Arabidopsis development. Similar inhibitory effects on plant development were observed in plants with inhibited pyrimidine *de novo* synthesis by PALA acting on aspartate transcarbamoylase (ATC) in line with ATC amiRNA lines (Bellin *et al*., 2021a, Bellin *et al*., 2023a, Slocum *et al*., 2023). In case of ATC and IMPDH mutants, reduced and most likely limiting nucleotide amounts were detected (Bellin *et al*., 2023b), Figure 7). We propose that such nucleotide limitation is fundamental for the observed developmental delay. It was suggested that in the embryos nucleotide reserves are depleted around 4 days after germination (Slocum *et al*., 2023). According to our analysis, we assume that this point is reached already 2 days after germination as in our mutants markedly reduced levels of GMP and in tendency also XMP and GTP and other nucleotides tested were seen at this time point (Figure 6, S9) before photosynthesis is fully established. This happens at a time point around 4 days after germination (Pipitone *et al*., 2021). The observation of decreased XMP levels but increased IMP levels, latter being the substrate of IMPDH indicated that in fact IMPDH represents the bottleneck in guanylate biosynthesis and furthermore affects adenylate and cytidylate amounts. Now the question arises about the mechanism which translates limiting nucleotides into developmental delays, going along with impaired photosynthesis, reduced chlorophyll levels and reduced biomass (Figure 3 and 4). In our transcriptome analysis “ribosome” due to reductions in mainly cytosolic ribosomal RNA levels showed up as major GO-Term (Figure 6) and reduced transcription/translation of photosynthesis related genes in plastid and cytosol can be seen as consequence (Figure 4D). It is meanwhile known that nucleotide limitation represses TOR activity, which then leads to reductions in ribosomal RNA (Busche *et al*., 2021). Among ribosomal genes that were reduced in expression RPS6A was identified, which is a target protein of S6K–TOR signaling. (Liu *et al*., 2025), underlining participation of TOR in IMPDH2-dependent seedling development. Reduced plastidic ribosomal RNA and DNA contents were observed in knock-down lines of *CTPS2*, another key enzyme in nucleotide synthesis (Alamdari *et al*., 2021, Bellin *et al*., 2021b). Homozygous crosses of *impdh1-1* with *impdh2-0* do not survive, however if only one functional *IMPDH1* allele is present, strongly growth retarded plants were identified, suffering from problems in plastid rRNA processing and thus ribosomal stress (Supplemental Table S1; Maekawa *et al*., 2024). In general, nucleotide limitations appear to act on (plastid) rRNA and DNA production in a direct way, simply because they are precursors but in addition via TOR-signaling. The observation of stronger effects on plastid RNA and DNA may be explained by the dominating role of chloroplasts in autotrophic tissues, which are mainly used in corresponding experiments. In those tissues chloroplasts can harbor most of the protein, DNA and RNA inventory of a cell.

As already pointed out above, Arabidopsis IMPDH2 is the dominant isoform with respect to its expression level which is higher than IMPDH1 in nearly all leaf and root tissues which are shown in the PLANT SINGLE CELL BROWSER (Ma *et al*. 2020) and tissues analyzed previously (Maekawa *et al*. 2024). In line with this observation, both *impdh2* knock-out lines demonstrate the highest impact of this isoform on plant development. Phenotypic alterations in *impdh2-0* and *2-1* appear early in development, not seen in wild type (WT) plants and only in a milder version in *IMPDH1* mutants but disappear during further plant development. These include chlorosis and reduced FW in 3 to 8 days old seedlings (Figure 3). The impact of IMPDH2 on early seedling development is further underlined by *IMPDH2* promotor-reporter studies where strongest reporter GUS staining was observed in 4-day-old seedlings (Fig. 7B). When these seedlings were grown on MS media, the supplementation with guanine or GMP reduced GUS staining, in contrast, the supply of the IMPDH inhibitor MPA led to a strong increase in promoter activity, indicating that guanylate availability regulates *IMPDH2* expression (Figure S8B). Moreover, GMP supplementation could rescue low chlorophyll amounts in *impdh2-0* mutants revealing that IMPDH is not only representing the bottleneck in guanylate synthesis but is also directly responsible for the observed impairment of photosynthesis (Figure S8A).

Besides reduced expression of ribosomal RNA encoding genes and photosynthesis related genes, genes related to nucleotide metabolism were altered in expression according to our transcriptome analysis (Figure 6F-H). Whereas genes involved in purine metabolism were mostly downregulated, pyrimidine metabolic genes were mostly upregulated. Strongest upregulation (1.73 log2 fold change impdh2-o vs WT) was seen for PUMPKIN (UMK5), involved in plastid RNA binding and splicing (Schmid *et al*., 2019). Another strongly upregulated gene was AT3G22450 (log_2_fold change 2.78) which encodes ribosomal protein uL18-L1. This protein is required for proper splicing of mitochondrial introns. Presumably in both cases *impdh2* mutants attempt to counteract impaired organellar RNA processing. Transporters involved in organellar energy balancing were also found upregulated (TAACL, APC1, NTT2) likely in response to reduced energy status. One gene, At3g20240, encoding the closest homolog to pANT1 (plastidial adenine nucleotide transporter) was found upregulated (Figure 6 G). It is tempting to speculate, that the encoded protein functions in IMP export from plastids (Haferkamp and Schmitz-Esser, 2012, Witte and Herde, 2020). If so, increased export of IMP to overcome the bottleneck of GMP synthesis in *impdh2* would lead to the observed IMP increase (in the cytosol) and thus remove substrate for AMP synthesis in plastids, Another possible explanation of reduced AMP levels is feedback inhibition of purine biosynthesis by IMP accumulating in plastids (Chen *et al*., 2022)

Arabidopsis harbors two IMPDH isoforms (Collart *et al*. 1996; Witte and Herde 2020) which share 85% identical amino acids. However, only one IMPDH isoform was found in about half of the multicellular species analyzed (Figure S1), suggesting that the presence of two or three isoforms in many other species must have a reason besides pure redundancy behind it. Vertebrates for example, harbor two isoforms and point mutations in either one of these lead to the manifestation of isoform specific diseases, indicating that each isoform exhibits individual functions (Burrell and Kollman 2022). When inspecting siliques, *impdh1-1* revealed 5% empty positions indicating aborted seeds and lower 1000 seed weight, compared to wild types, suggesting a prominent role of this isoform in embryo and seed development. This hypothesis is further substantiated by phenotypic alterations of approximately 10% of *IMPDH1* ox#2 and ox#3 seedlings, which exhibit 3-4 cotyledons of various morphologies (Figure 5 A). Moreover, transcript levels of auxin markers, most markedly *SAUR* (small auxin upregulated) 9 and 10 (Figure 5C) allow to speculate about elevated auxin levels in *IMPDH1* ox#2 embryos, while *impdh1-1* embryos presumably exhibited contrasting effects (Figure 5C). As we observed increased expression of the apical marker *WOX2* but decreased expression of the basal marker *WOX8* a shift in auxin levels towards the apex can be assumed. PIN1 is crucial for auxin transport within the apical to basal tissue and reduced *PIN1* levels were observed in the *IMPDH1* ox#2, explaining shifted auxin amounts. *PIN1* knock-out mutants exhibited the same cotyledon phenotype as those in the *IMPDH1* ox seedlings in 12% of corresponding seedlings (Robert *et al*., 2015). This finding underscores the importance of functional IMPDH1 levels for optimal embryo development, presumably by assuring auxin polarization within the developing embryo. As Auxin polarization in the apical tissue of embryo leads to formation of cotyledons, an excessive amount of auxin in the apical part of embryo can lead to changes in the number of auxin maxima resulting in changes of cotyledon number (Huang *et al*., 2010). We think that IMPDH isoforms in vertebrates are functionally separated, whereas in Arabidopsis the two isoforms show higher redundancy but still exhibit partially individual roles explained by their expression profiles.

IMPDH and CTPS, the latter enzyme catalyzing CTP synthesis from UTP, were found together in macromolecular structures in Drosophila and human cell lines, forming cytoophidia or rods and ring structures (Keppeke *et al*., 2015, Liu, 2016). We could obtain indications for the capacity of CTPS isoforms 3-5 to form cytoophidia in previous studies (Daumann *et al*., 2018). Furthermore, rod and ring structures were also seen for CFP-IMPDH1 and GFP-IMPDH2 after transient expression in *Nicotiana benthamiana* (Figure S3), allowing us to speculate about common macromolecular structures for both enzymes. We chose to analyze this with the expression of various combinations of BIFC constructs and observed positive results for interactions with IMPDH1 and 2, and IMPDHs with CTPS isoforms. IMPDH is the rate-limiting enzyme for GTP *de novo* synthesis, and GTP has been shown to act as essential activator of CTPS in the biosynthesis of CTP. We think this interaction between purine and pyrimidine metabolism is critical for the maintenance of nucleotide homeostasis.

Formation of macromolecular structures represents another layer of regulation as it was observed that human CTPS can polymerize in response to altered nucleotide levels and by this strongly affected the activity of the corresponding enzymes (Lynch *et al*., 2017, Lynch and Kollman, 2020). However, filament formation can have many diverse functions in a cell apart from activity regulation, by this affecting protein stability, cell structure, cell adhesion and cell to cell communication (Guo and Liu, 2023) In addition, TOR was shown to mediate CTPS-filament assembly in Drosophila (Sun and Liu, 2019). In leaf discs expressing *IMPDH2:GFP*, increased formation of macromolecular structures was observed after ATP (0.5 mM) supplementation (Figure S5). As the presence of ATP indicates a high energy status which should promote anabolic pathways, we can assume that IMPDH macromolecular structures resemble the active enzyme conformation.

## Experimental procedures

### Plant growth

WT and transgenic *Arabidopsis thaliana* (L.) Heynh. Plants (ecotype Columbia) were used for DNA isolation, tissue collection, and phenotypic inspection throughout. Standardized ED73 soil with 10% sand (Einheitserde und Humuswerke Patzer) or ½ MS agar plates were used for plant growth under short day (SD) conditions with 10 h light (120 μmol quanta m^−2^s^−1^); 22°C and 60% humidity. LED lights (Valoya NS1, Valoya, Finland) were used for illumination. For growth experiments on sterile agar plates and liquid cultures, seeds were surface sterilized and placed on half-strength MS supplemented with 0.1% (w/v) sucrose. For supplementation experiments, the medium was enriched with either 0.5 mM guanine, 0.5 mM GMP, or 10 µM mycophenolic acid (MPA). Seeds were incubated for 48 h in the dark at 4°C for imbibition (Weigel and Glazebrook, 2002). Liquid cultivation was performed in 6 well plates with 5 ml media per well and constant rotation at 100 rpm.

### Construction of IMPDH2 reporter lines, *IMPDH1* complementation lines and *IMPDH1/2* overexpression lines

Construction of promoter GUS lines by gateway cloning into the pBGWF7.0 vector was performed as described before (Hickl *et al*., 2021). For this, 1889 bp upstream of the start codon of At1g16350 (*IMPDH2*) were amplified from WT gDNA and inserted into pBGWF7.0. Primers used are given in Table S2. pMDC123 was used for complementation of *impdh1-1* with the *IMPDH1* full length native genomic construct including promoter as given above (Curtis and Grossniklaus, 2003). Overexpression of both *IMPDH1* and *IMPDH2* from WT cDNA was achieved by Gateway Cloning into pUBC (Grefen *et al*., 2010) resulting in control of transgene expression by the ubiquitin10 promoter. Correctness of the cloning procedure was checked by enzyme digestion and Sanger sequencing (Eurofins, Ebersberg). Transformation into WT plants and *IMPDH1* mutants was performed according to the floral inoculation procedure (Narusaka *et al*., 2010)

### Transcript quantification with qRT PCR

The isolation of RNA, subsequent preparation of cDNA, and PCR were conducted in accordance with the methods outlined in (Ohler *et al*., 2019). Leaf tissue was frozen using liquid nitrogen, followed by grinding to a fine powder. Subsequently, RNA was isolated using the Nucleospin RNA Plant Kit (Macherey-Nagel, Düren, Germany), in accordance with the manufacturer’s instructions. The isolated RNA was then transcribed into cDNA using the qScript cDNA Synthesis Kit (Quantabio, United States). The Primers used are listed in Supplemental Table S2.

### Transcriptome analysis

A comprehensive expression analysis of WT and impdh2 knockout plants was performed to obtain an overview of the transcriptome during early seedling development. For this purpose, WT, *impdh2-0* and *ami-ctps2-2* seedlings at the same stage of development were harvested in the chlorotic growth phase and in the greening phase. Two hours after the start of the light phase, the seedlings without roots were harvested and transferred directly to liquid nitrogen. Isolated RNA samples were sent to Novogene (China), which created the RNA library and performed transcriptome sequencing. The online tool REVIGO (Supek *et al*., 2011) was used to evaluate the GO term analysis.

### Protein extraction, gel-electrophoresis and immunoblotting

50-100 mg leaf samples were grinded in liquid nitrogen and subsequently extracted with protein extraction buffer (50 mM Hepes-KOH, pH 7.8, 5 mM MgCl, 1 mM ethylenediaminetetraacetic acid, 2 mM dithiothreitol, 0.01 % DDM and 2% (v/v) protease inhibitor cocktail (Sigma, St Louis, MO). Supernatants were then centrifuged at 13,000 *g* for 1 min at 4 °C. Protein extracts were supplemented with 4× Laemmli buffer and the samples were stored at -80°C until use. SDS PAGE was performed with precast SDS gels (BioRad, Hercules, CA, USA) according to the manufacturers protocol.

Immunoblots were essentially performed as given in (Ohler *et al*., 2019), here the proteins were separated on SDS-gels and transferred onto a nitrocellulose membrane by the semi-dry Transblot Turbo Transfer System (Bio-Rad, Hercules, CA, USA). Primary antibodies were obtained from Agrisera (Vännäs, Sweden).

### Photosynthetic activity was measured using pulse-amplitude-modulation (PAM)

Chlorophyll fluorescence was measured using a MINI Imaging pulse-amplitude modulation fluorometer (Walz, www.walz.com). Prior to measurement, plants were adapted to the dark for 10 min. The quantum yield of photochemical energy conversion on PSII Φ_(PSII)_and the quantum yield of non-photochemical quenching of chlorophyll fluorescence Φ_(NPQ)_ and unregulated energy dissipation Φ_(NO)_ were determined (Kramer *et al*., 2004). Induction curves were run for 5.4 min and actinic light (76 μmol photons m^−2^ s^−1^) provided from 40 seconds time point onwards, every 20 seconds.

### Statistical analysis

Statistical analyses of numerical data were performed using GraphPad Prism 10. To test for significant differences between two groups, a one-way ANOVA test followed by a Dunnett multiple comparison test was performed (p-value < 0.05 = *; p-value < 0.01 = **, p-value < 0.001 = ***, p-value < 0.0001 = ****). The ‘Rapid Publication-Ready MS Word Tables Using Two-Way ANOVA’ software (Assaad *et al*., 2015) (https://houssein-assaad.shinyapps.io/TwoWayANOVA/) was used for a letter-based representation of the pairwise comparisons, using a two-sided ANOVA followed by post-hoc Tukey HSD testing

### Transient expression in *Nicothiana benthamiana* and Microscopy

Transient expression of IMPDH fused to YFP, GFP or CFP was performed as detailed in Walter *et al*. (2004). Therefore, six weeks old *N. benthamiana* leaves were infiltrated through the lower epidermis. After four to five days leaves were analyzed for the presence of fluorescence signals with a Leica TCS SP5II microscope (CFP 355 nm excitation and 495–530 nm detection YFP 514 nm excitation and 525–582 nm detection, GFP 488 nm excitation and 505-540 nm detection of emission through a HCX PL APO 63 × 1.2 W water immersion objective). Chlorophyll autofluorescence was detected with 514 nm excitation and a 651–704 nm emission wavelength.

### Chlorophyll quantification

Leaf tissue was flash frozen in liquid nitrogen and subsequently ground to a fine powder. Subsequently, 80% ethanol was added, and the sample was subjected to heating at a temperature of 95°C for 10min followed by sedimentation of the insoluble contents by centrifugation at 20,000 g for 10min. The chlorophyll content of the sample was determined by measuring its absorbance at 652 nm and calculating Chlorophyll A and B (Porra and Scheer, 2019)

### Lipid extraction

Seeds (100mg) were ground in liquid nitrogen and lipids were extracted by addition of 750 µl isopropanol under constant shaking for 12h at 4°C (Reiser *et al*., 2004). Lipid extracts were then dried and dissolved in chloroform and subsequently purified using solid-phase extraction (SPE). In the thin-layer chromatography (TLC) procedure, lipid extract and standards (rice oil or triacylglycerol) were applied to silica gel plates and developed in hexane diethyl ether acetic acid (70:30:1). The presence of lipid bands was determined with iodine staining.

### Nucleotide quantification

For cultivation of 0, 12 and 48 hours seed and seedling samples a previously described protocol was used (Niehaus *et al*., 2022). Eight-day old seedlings were grown on soil as described above. Samples for 0 and 12 hours were lyophilized prior to nucleotide extraction to remove solid frozen liquid interfering with sample grinding. Nucleotide quantification was then performed as described in (Straube *et al*., 2021) with modifications from (Straube *et al*., 2023).

## Acknowledgements

T.M. conceived and supervised the study, obtained funding, and provided resources. E.D. generated all mutants and performed characterization, and phenotyping. E.D.,T.M. performed analysis and interpretation of transcriptomic and metabolomic data. L.F, M.H. performed metabolomic data analysis. M.H., C.P.W advised and interpreted metabolite quantification.

S.Z. supported *E. coli* complementation assays. T.M. and E.D. wrote the original draft. All authors reviewed and agreed to the final manuscript.

We thank Nicole Frankenberg-Dinkel for providing strains from the Keio Collection and Leo Bellin for fruitful discussions throughout the work.

## Data Availability statement

“The data discussed in this publication have been deposited in NCBI’s Gene Expression Omnibus (Edgar *et al*., 2002) and are accessible through GEO Series accession number GSE312534 (https://www.ncbi.nlm.nih.gov/geo/query/acc.cgi?acc=GSE268410)

## References

Alamdari, K., Fisher, K.E., Tano, D.W., Rai, S., Palos, K., Nelson, A.D.L. and Woodson, J.D. (2021) Chloroplast quality control pathways are dependent on plastid DNA synthesis and nucleotides provided by cytidine triphosphate synthase two. New Phytol, 231, 1431–1448.

Assaad, H.I., Hou, Y., Zhou, L., Carroll, R.J. and Wu, G. (2015) Rapid publication-ready MS-Word tables for two-way ANOVA. Springerplus, 4, 33.

Bellin, L., Del Cano-Ochoa, F., Velazquez-Campoy, A., Möhlmann, T. and Ramon-Maiques, S. (2021a) Mechanisms of feedback inhibition and sequential firing of active sites in plant aspartate transcarbamoylase. Nat Commun, 12, 947.

Bellin, L., Garza Amaya, D.L., Scherer, V., Pruss, T., John, A., Richter, A. and Möhlmann, T. (2023a) Nucleotide imbalance, provoked by downregulation of Aspartate Transcarbamoylase impairs cold acclimation in Arabidopsis. Molecules, 28.

Bellin, L., Melzer, M., Hilo, A., Garza Amaya, D.L., Keller, I., Meurer, J. and Mohlmann, T. (2023b) Nucleotide limitation results in impaired photosynthesis, reduced growth and seed yield together with massively altered gene expression. Plant Cell Physiol, 64, 1494–1510.

Bellin, L., Scherer, V., Dörfer, E., Lau, A., Vicente, A.M., Meurer, J., Hickl, D. and Möhlmann, T. (2021b) Cytosolic CTP production limits the establishment of photosynthesis in Arabidopsis. Front Plant Sci, 12, 789189.

Bowne, S.J., Sullivan, L.S., Blanton, S.H., Cepko, C.L., Blackshaw, S., Birch, D.G., Hughbanks-Wheaton, D., Heckenlively, J.R. and Daiger, S.P. (2002) Mutations in the inosine monophosphate dehydrogenase 1 gene (IMPDH1) cause the RP10 form of autosomal dominant retinitis pigmentosa. Hum Mol Genet, 11, 559–568.

Buey, R.M., Ledesma-Amaro, R., Velazquez-Campoy, A., Balsera, M., Chagoyen, M., de Pereda, J.M. and Revuelta, J.L. (2015) Guanine nucleotide binding to the Bateman domain mediates the allosteric inhibition of eukaryotic IMP dehydrogenases. Nat Commun, 6, 8923.

Burrell, A.L. and Kollman, J.M. (2022) IMPDH dysregulation in disease: a mini review. Biochem Soc Trans, 50, 71–82.

Busche, M., Scarpin, M.R., Hnasko, R. and Brunkard, J.O. (2021) TOR coordinates nucleotide availability with ribosome biogenesis in plants. Plant Cell.

Cao, Y. and Schubert, K.R. (2001) Molecular cloning and characterization of a cDNA encoding soybean nodule IMP dehydrogenase. Biochim Biophys Acta, 1520, 242–246.

Carcamo, W.C., Satoh, M., Kasahara, H., Terada, N., Hamazaki, T., Chan, J.Y., Yao, B., Tamayo, S., Covini, G., von Muhlen, C.A. and Chan, E.K. (2011) Induction of cytoplasmic rods and rings structures by inhibition of the CTP and GTP synthetic pathway in mammalian cells. PLoS One, 6, e29690.

Carr, S.F., Papp, E., Wu, J.C. and Natsumeda, Y. (1993) Characterization of human type I and type II IMP dehydrogenases. J Biol Chem, 268, 27286–27290.

Chang, C.C., Keppeke, G.D., Sung, L.Y. and Liu, J.L. (2018) Interfilament interaction between IMPDH and CTPS cytoophidia. FEBS J, 285, 3753–3768.

Chang, Y.F. and Carman, G.M. (2008) CTP synthetase and its role in phospholipid synthesis in the yeast Saccharomyces cerevisiae. Prog Lipid Res, 47, 333–339.

Chen, X., Kim, S.H., Rhee, S. and Witte, C.P. (2022) A plastid nucleoside kinase is involved in inosine salvage and control of purine nucleotide biosynthesis. Plant Cell.

Collart, F.R., Osipiuk, J., Trent, J., Olsen, G.J. and Huberman, E. (1996) Cloning and characterization of the gene encoding IMP dehydrogenase from *Arabidopsis thaliana*. Gene, 174, 217–220.

Curtis, M.D. and Grossniklaus, U. (2003) A gateway cloning vector set for high-throughput functional analysis of genes in planta. Plant Physiol, 133, 462–469.

Daumann, M., Hickl, D., Zimmer, D., DeTar, R.A., Kunz, H.H. and Möhlmann, T. (2018) Characterization of filament-forming CTP synthases from *Arabidopsis thaliana*. Plant J, 96, 316–328.

Edgar, R., Domrachev, M. and Lash, A.E. (2002) Gene Expression Omnibus: NCBI gene expression and hybridization array data repository. Nucleic Acids Res, 30, 207–210.

Grefen, C., Donald, N., Hashimoto, K., Kudla, J., Schumacher, K. and Blatt, M.R. (2010) A ubiquitin-10 promoter-based vector set for fluorescent protein tagging facilitates temporal stability and native protein distribution in transient and stable expression studies. Plant J, 64, 355–365.

Guo, C.J. and Liu, J.L. (2023) Cytoophidia and filaments: you must unlearn what you have learned. Biochem Soc Trans, 51, 1245–1256.

Haferkamp, I. and Schmitz-Esser, S. (2012) The plant mitochondrial carrier family: functional and evolutionary aspects. Front Plant Sci, 3, 2.

Hickl, D., Scheuring, D. and Möhlmann, T. (2021) CTP Synthase 2 from *Arabidopsis thaliana* is required for complete embryo development. Front Plant Sci, 12, 652434.

Holzl, G. and Dormann, P. (2019) Chloroplast lipids and their biosynthesis. Annu Rev Plant Biol, 70, 51–81.

Jackson, R.C., Weber, G. and Morris, H.P. (1975) IMP dehydrogenase, an enzyme linked with proliferation and malignancy. Nature, 256, 331–333.

Johnson, M.C. and Kollman, J.M. (2020) Cryo-EM structures demonstrate human IMPDH2 filament assembly tunes allosteric regulation. Elife, 9.

Keppeke, G.D., Calise, S.J., Chan, E.K. and Andrade, L.E. (2015) Assembly of IMPDH2-based, CTPS-based, and mixed rod/ring structures is dependent on cell type and conditions of induction. J Genet Genomics, 42, 287–299.

Keppeke, G.D., Chang, C.C., Peng, M., Chen, L.Y., Lin, W.C., Pai, L.M., Andrade, L.E.C., Sung, L.Y. and Liu, J.L. (2018) IMP/GTP balance modulates cytoophidium assembly and IMPDH activity. Cell Div, 13, 5.

Kim, Y.K., Kim, S., Shin, Y.J., Hur, Y.S., Kim, W.Y., Lee, M.S., Cheon, C.I. and Verma, D.P. (2014) Ribosomal protein S6, a target of rapamycin, is involved in the regulation of rRNA genes by possible epigenetic changes in Arabidopsis. J Biol Chem, 289, 3901–3912.

Kramer, D.M., Johnson, G., Kiirats, O. and Edwards, G.E. (2004) New fluorescence parameters for the determination of QA redox state and excitation energy fluxes. Photosynth Res, 79, 209.

Krämer, M., Dorfer, E., Hickl, D., Bellin, L., Scherer, V. and Möhlmann, T. (2022) Cytidine Triphosphate Synthase Four from *Arabidopsis thaliana* attenuates drought stress effects. Front Plant Sci, 13, 842156.

Leroch, M., Kirchberger, S., Haferkamp, I., Wahl, M., Neuhaus, H.E. and Tjaden, J. (2005) Identification and characterization of a novel plastidic adenine nucleotide uniporter from *Solanum tuberosum*. J. Biol Chem, 280, 17992–18000.

Levitzki, A. and Koshland, D.E., Jr. (1972) Role of an allosteric effector. Guanosine triphosphate activation in cytosine triphosphate synthetase. Biochemistry, 11, 241–246.

Liu, J.L. (2016) The Cytoophidium and Its Kind: Filamentation and Compartmentation of Metabolic Enzymes. Annu Rev Cell Dev Biol, 32, 349–372.

Liu, Y., Hu, J., Duan, X., Ding, W., Xu, M. and Xiong, Y. (2025) Target of Rapamycin (TOR): A master regulator in plant growth, development, and stress responses. Annu Rev Plant Biol, 76, 341–371.

Lunn, F.A., MacDonnell, J.E. and Bearne, S.L. (2008) Structural requirements for the activation of Escherichia coli CTP synthase by the allosteric effector GTP are stringent, but requirements for inhibition are lax. J. Biol Chem, 283, 2010–2020.

Lynch, E.M., Hicks, D.R., Shepherd, M., Endrizzi, J.A., Maker, A., Hansen, J.M., Barry, R.M., Gitai, Z., Baldwin, E.P. and Kollman, J.M. (2017) Human CTP synthase filament structure reveals the active enzyme conformation. Nat Struct Mol Biol, 24, 507–514.

Lynch, E.M. and Kollman, J.M. (2020) Coupled structural transitions enable highly cooperative regulation of human CTPS2 filaments. Nat Struct Mol Biol, 27, 42–48.

Maekawa, S., Nishikawa, I. and Horiguchi, G. (2024) Impaired inosine monophosphate dehydrogenase leads to plant-specific ribosomal stress responses in *Arabidopsis thaliana*. J Plant Res, 137, 1091–1104.

Maraschin Fdos, S., Memelink, J. and Offringa, R. (2009) Auxin-induced, SCF(TIR1)-mediated poly-ubiquitination marks AUX/IAA proteins for degradation. Plant J, 59, 100–109.

Moffatt, B.A. and Ashihara, H. (2002) Purine and pyrimidine nucleotide synthesis and metabolism, pp. 1–20.

Narusaka, M., Shiraishi, T., Iwabuchi, M. and Narusaka, Y. (2010) The floral inoculating protocol: a simplified Arabidopsis thaliana transformation method modified from floral dipping. Plant Biotechnol-Nar, 27, 349–351.

Niehaus, M., Straube, H., Specht, A., Baccolini, C., Witte, C.P. and Herde, M. (2022) The nucleotide metabolome of germinating *Arabidopsis thaliana* seeds reveals a central role for thymidine phosphorylation in chloroplast development. Plant Cell, 34, 3790–3813.

Ohler, L., Niopek-Witz, S., Mainguet, S.E. and Möhlmann, T. (2019) Pyrimidine Salvage: Physiological Functions and Interaction with Chloroplast Biogenesis. Plant Physiol, 180, 1816–1828.

Pipitone, R., Eicke, S., Pfister, B., Glauser, G., Falconet, D., Uwizeye, C., Pralon, T., Zeeman, S.C., Kessler, F. and Demarsy, E. (2021) A multifaceted analysis reveals two distinct phases of chloroplast biogenesis during de-etiolation in Arabidopsis. Elife, 10.

Porra, R.J. and Scheer, H. (2019) Towards a more accurate future for chlorophyll a and b determinations: the inaccuracies of Daniel Arnon’s assay. Photosynth Res, 140, 215–219.

Reiser, J., Linka, N., Lemke, L., Jeblick, W. and Neuhaus, H.E. (2004) Molecular physiological analysis of the two plastidic ATP/ADP transporters from Arabidopsis. Plant Physiol, 136, 3524–3536.

Robert, H.S., Grunewald, W., Sauer, M., Cannoot, B., Soriano, M., Swarup, R., Weijers, D., Bennett, M., Boutilier, K. and Friml, J. (2015) Plant embryogenesis requires AUX/LAX-mediated auxin influx. Development, 142, 702–711.

Schmid, L.M., Ohler, L., Möhlmann, T., Brachmann, A., Muino, J.M., Leister, D., Meurer, J. and Manavski, N. (2019) PUMPKIN, the sole plastid UMP Kinase, associates with group II introns and alters their metabolism. Plant Physiol, 179, 248–264.

Sifri, C.D., Wilson, K., Smolik, S., Forte, M. and Ullman, B. (1994) Cloning and sequence analysis of a Drosophila melanogaster cDNA encoding IMP dehydrogenase. Biochim Biophys Acta, 1217, 103–106.

Slocum, R.D., Peña, C.M. and Liu, Z.C. (2023) Transcriptional reprogramming of nucleotide metabolism in response to altered pyrimidine availability in Arabidopsis seedlings. Frontiers in Plant Science, 14.

Straube, H., Niehaus, M., Zwittian, S., Witte, C.P. and Herde, M. (2021) Enhanced nucleotide analysis enables the quantification of deoxynucleotides in plants and algae revealing connections between nucleoside and deoxynucleoside metabolism. Plant Cell, 33, 270–289.

Straube, H., Straube, J., Rinne, J., Fischer, L., Niehaus, M., Witte, C.P. and Herde, M. (2023) An inosine triphosphate pyrophosphatase safeguards plant nucleic acids from aberrant purine nucleotides. New Phytol, 237, 1759–1775.

Sun, Z. and Liu, J.L. (2019) mTOR-S6K1 pathway mediates cytoophidium assembly. J Genet Genomics, 46, 65–74.

Supek, F., Bosnjak, M., Skunca, N. and Smuc, T. (2011) REVIGO summarizes and visualizes long lists of gene ontology terms. PLoS One, 6, e21800.

Wang, C., Fourdin, R., Quadrado, M., Dargel-Graffin, C., Tolleter, D., Macherel, D. and Mireau, H. (2020) Rerouting of ribosomal proteins into splicing in plant organelles. Proc Natl Acad Sci U S A, 117, 29979–29987.

Weigel, D. and Glazebrook, J. (2002) Arabidopsis. A Laboratory Manual New York: Cold Spring Harbor Laboratory Press.

Witte, C.P. and Herde, M. (2020) Nucleotide Metabolism in Plants. Plant Physiol, 182, 63–78.

Witz, S., Jung, B., Fürst, S. and Möhlmann, T. (2012) De novo pyrimidine nucleotide synthesis mainly occurs outside of plastids, but a previously undiscovered nucleobase importer provides substrates for the essential salvage pathway in Arabidopsis. Plant Cell, 24, 1549–1559.

Zrenner, R., Stitt, M., Sonnewald, U. and Boldt, R. (2006) Pyrimidine and purine biosynthesis and degradation in plants. Annu. Rev. Plant Biol, 57, 805–836.

