## Supplemental figures for "Inosine Monophosphate Dehydrogenases are key players for functional development in Arabidopsis"

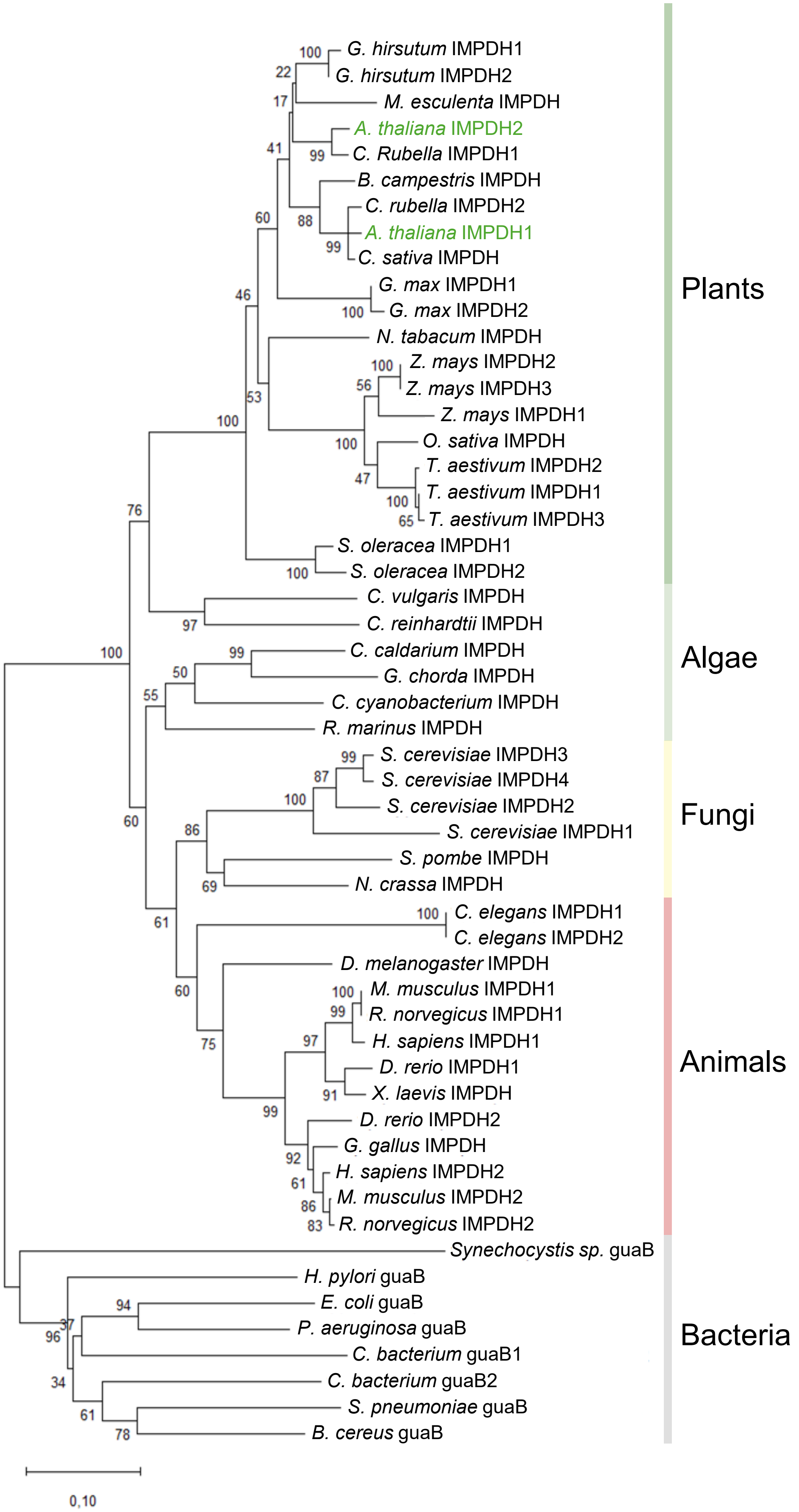


**Figure S1***:* **Phylogenetic tree of IMPDH isoforms across the kingdoms of life.** Maximum likelyhood, unrooted phylogenetic tree of selected members IMPDH family across all kingdoms of life. The analysis was performed with MEGA12.

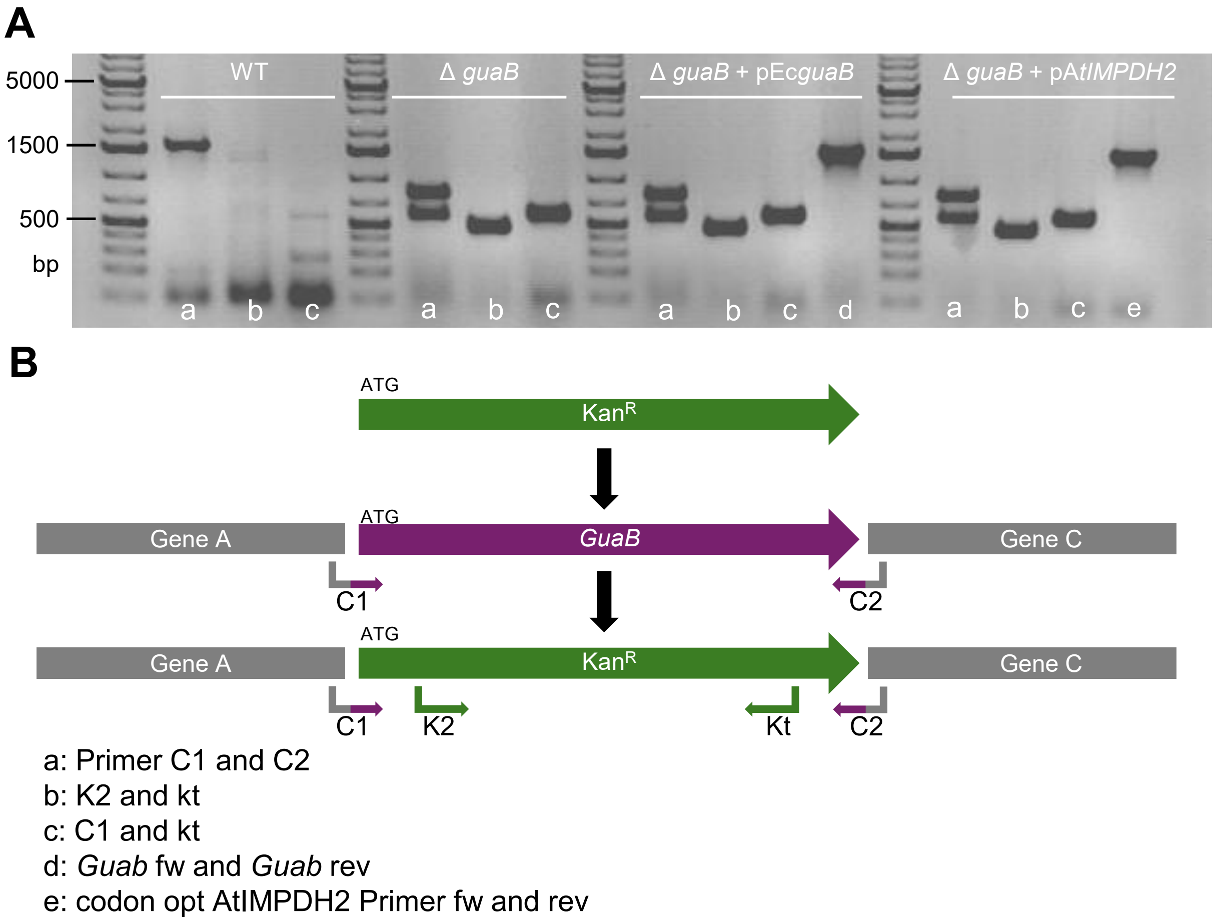


**Figure S2: Genotyping *E. coli IMPDH* complementation.**
PCR results of *E. coli* BW25113 (WT), Δ*guaB*, Δ*guaB* complemented with native *E.coli IMPDH* (*guaB*), and Δ*guaB* complemented with a for *E. coli* codon optimized version of At*IMPDH2*. Following primer combinations were used: (a) C1 and C2 which bind on 5‘ side of vector and gene start as well as gene end and 3‘ side of the vector , (b) K2 and Kt which bind on the kanamycin resistance gene in the Δ guaB strain, (c) C1 and Kt which bind on 5‘ side of vector and gene start and on the kanamycin resistance gene, (d) *guaB* gene specific, (e) *IMPDH2* gene specific. The used marker is 1 kb plus DNA-ladder from Thermo-Fisher.

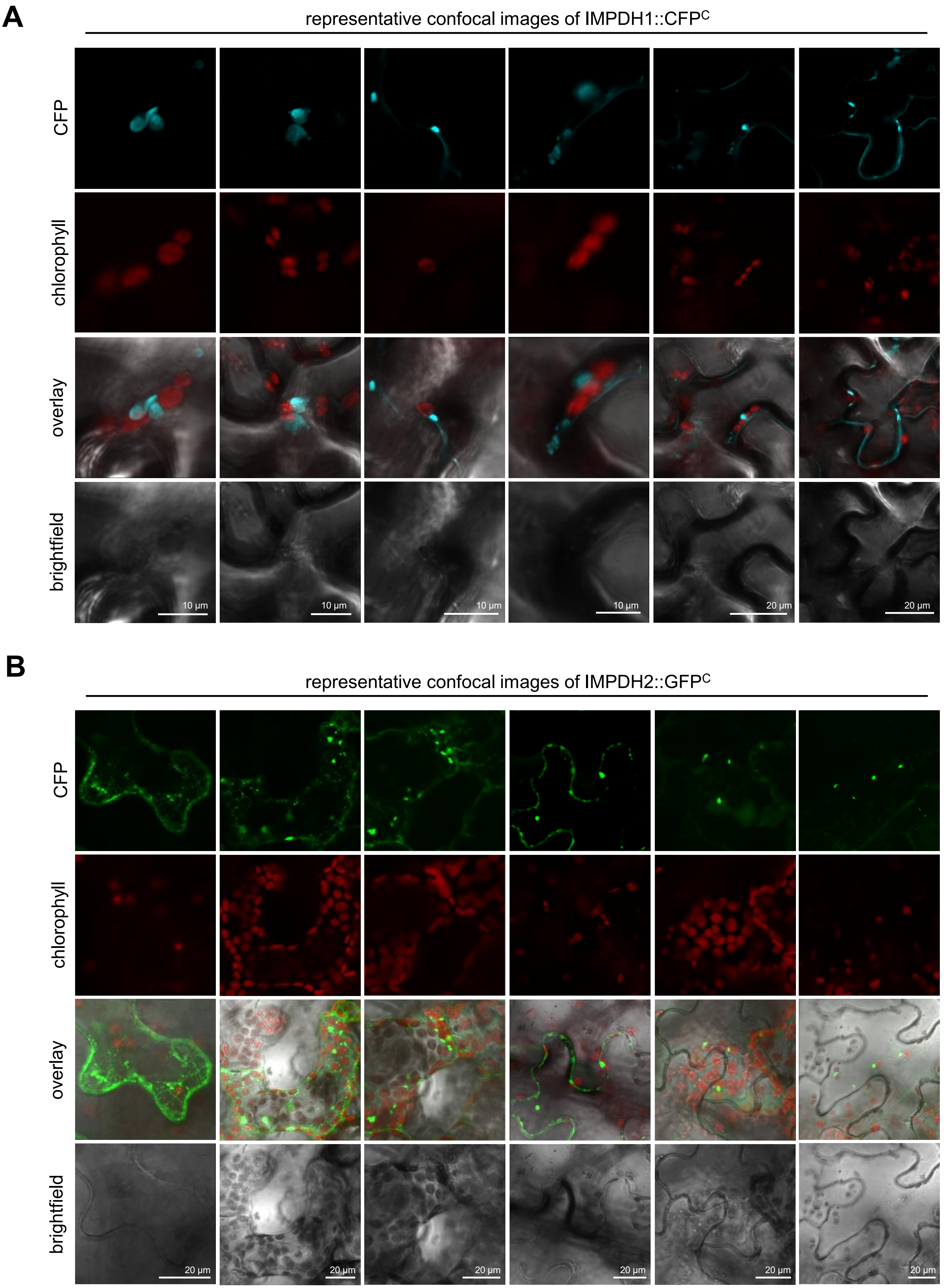


**Figure S3**: **Close ups of IMPDH Polymers.** (**A**) Detailed pictures of IMPDH1::CFP and (**B**) IMPDH2::GFP transiently expressed in *N. benthamiana*.

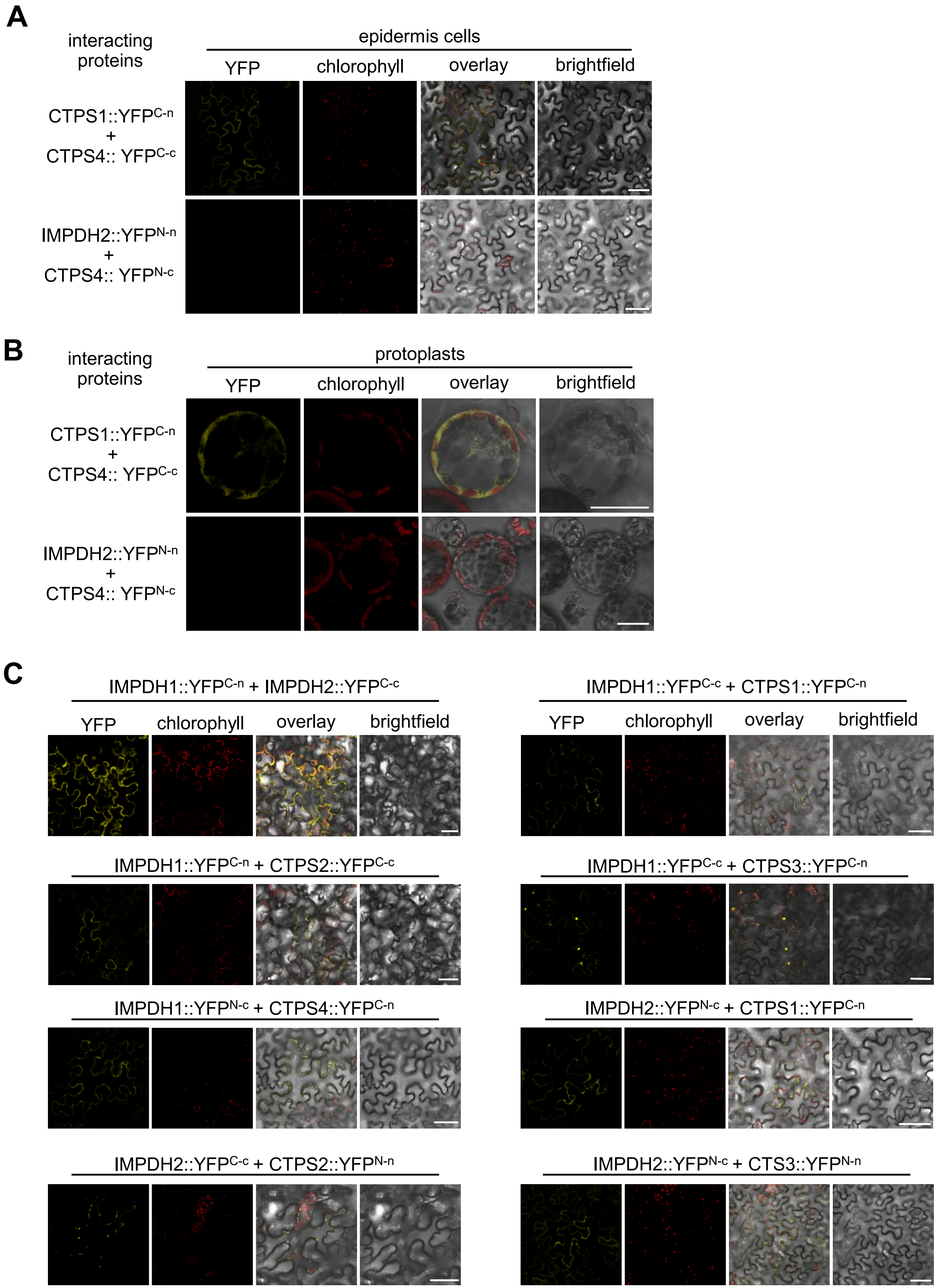


**Fig S4: BiFC analaysis of IMPDH isoforms and CTPS isoforms.** Constructs of full-length coding sequences of IMPDH and CTPS isoforms were fused upstream to the C- and N-terminus of YFP and transiently expressed in *N. benthamiana* leaves by agroinfiltration. YFP and chlorophyll fluorescence signals were recorded 4 d after infiltration in isolated protoplasts by confocal microscopy. Scalebars = 50 µm for epidermis pictures, and 20 µm for Protoplasts.

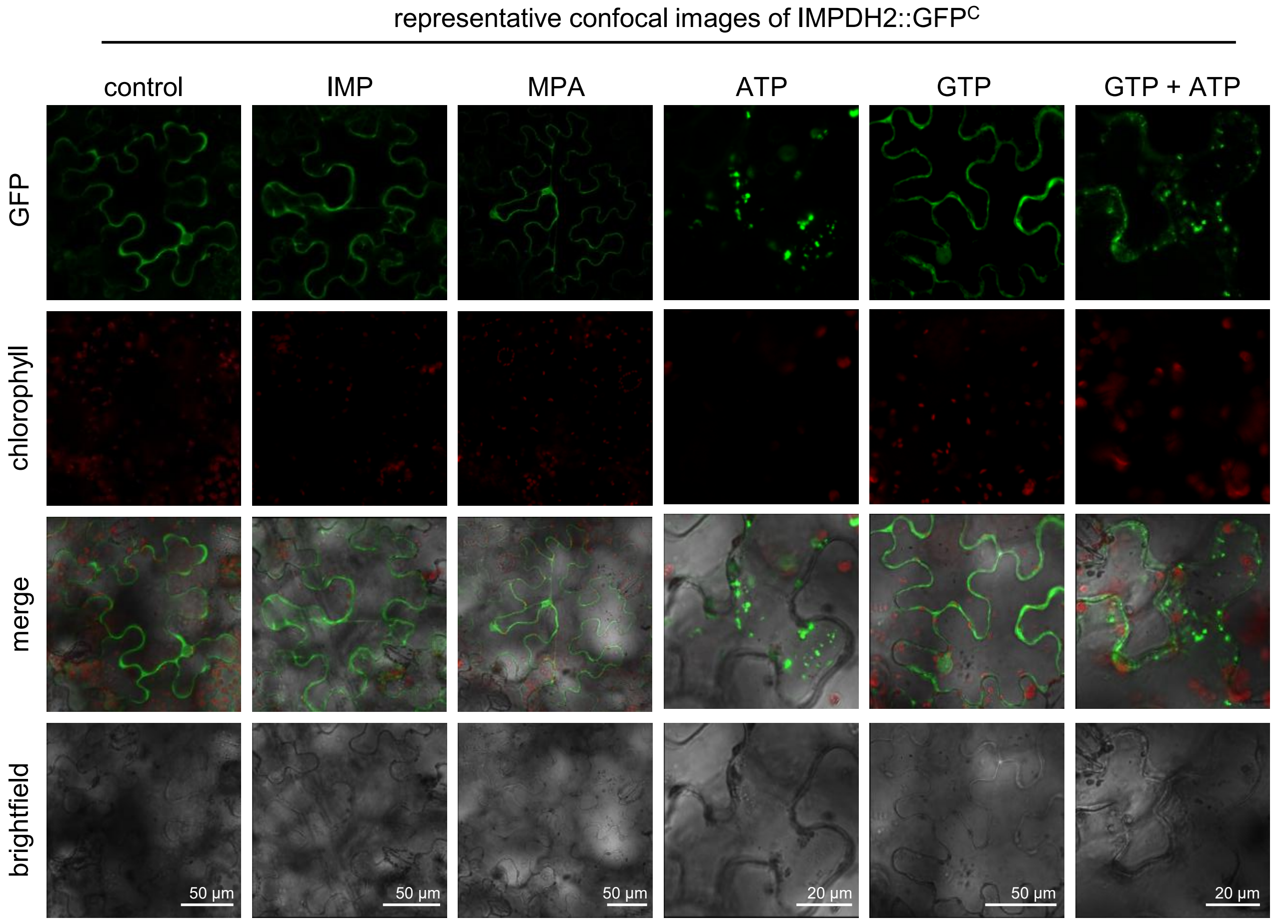


**Figure S5: Leaf disc experiment of IMPDH2::GFP.** Tobacco epidermis cells infiltrated with IMPDH2-GFP contruct. Leaf discs were cut out and incubated in 10mM MES-KOH pH 5.7 with 10 mM MgCl2 and 0.1% Triton X 100, supplemented with either 0.5 mM IMP; 10 µM MPA, 0.5 mM GTP; 0.5 mM ATP and 0.5 mM GTP + 0.5 mM ATP.

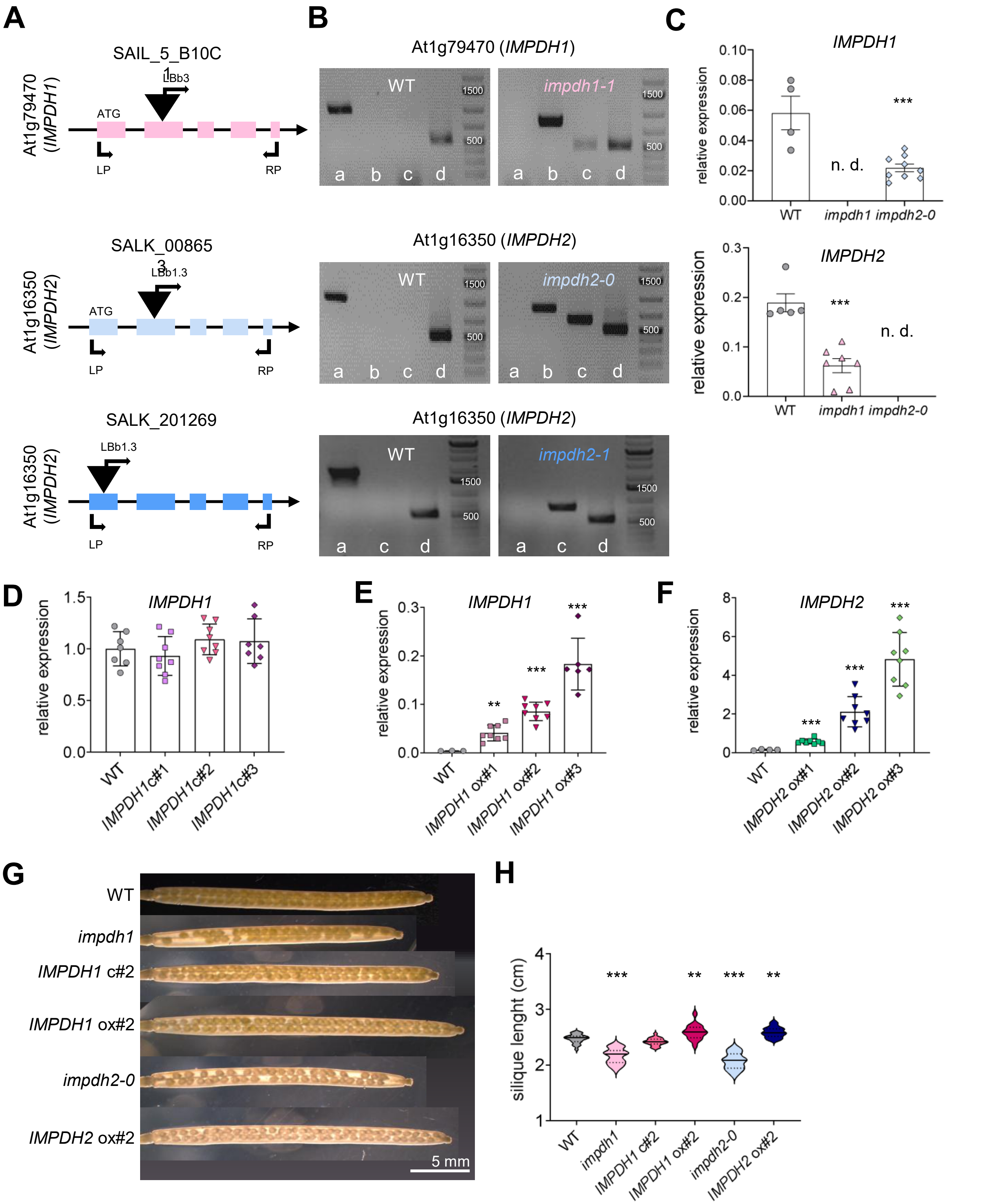


**Figure S6. Genotyping of *IMPDH1* and *IMPDH2* t-DNA Insertion, complementation and overexpression lines.** (**A**) Schematic drawing showing the position of t-DNA insertions based on the flanking sequences. Black line and colored boxes represent introns and exons. (**B**) Genotyping PCR of the t-DNA insertion knock-out lines. Genomic DNA was used for PCR reactions with the following primer combinations: (a) gene specific LP + RP, t-DNA-specific LBb3 or LBb1.3 and gene-specific (b) RP (c) LP-Primer and (d) *Ef1a* as a positive control. Line name equivalents: *impdh1* = SAIL_5_B10C1, *impdh2-0* = SALK_008653, and *impdh2-* = SALK_201269. (**C-F**) The relative expression of *IMPDH1* and *IMPDH2* transcript in 21 days old plants was determined by qRT-PCR. *Actin2* was used as a reference gene. Plants were raised under short day conditions (10h light and 14h dark. Microscopic inspection of seeds in developing siliques (**G**) Representative microscopic inspection of seeds in developing siliques and (**H**) silique length of mature siliques (n=20). Error bars represent ± SD. Asterisks indicate significant differences between WT and mutant using a one-way ANOVA followed by a dunnett's multiple comparisons test : * p-value ≤ 0.05, ** p-value ≤ 0.01, ***p-value ≤ 0.001.

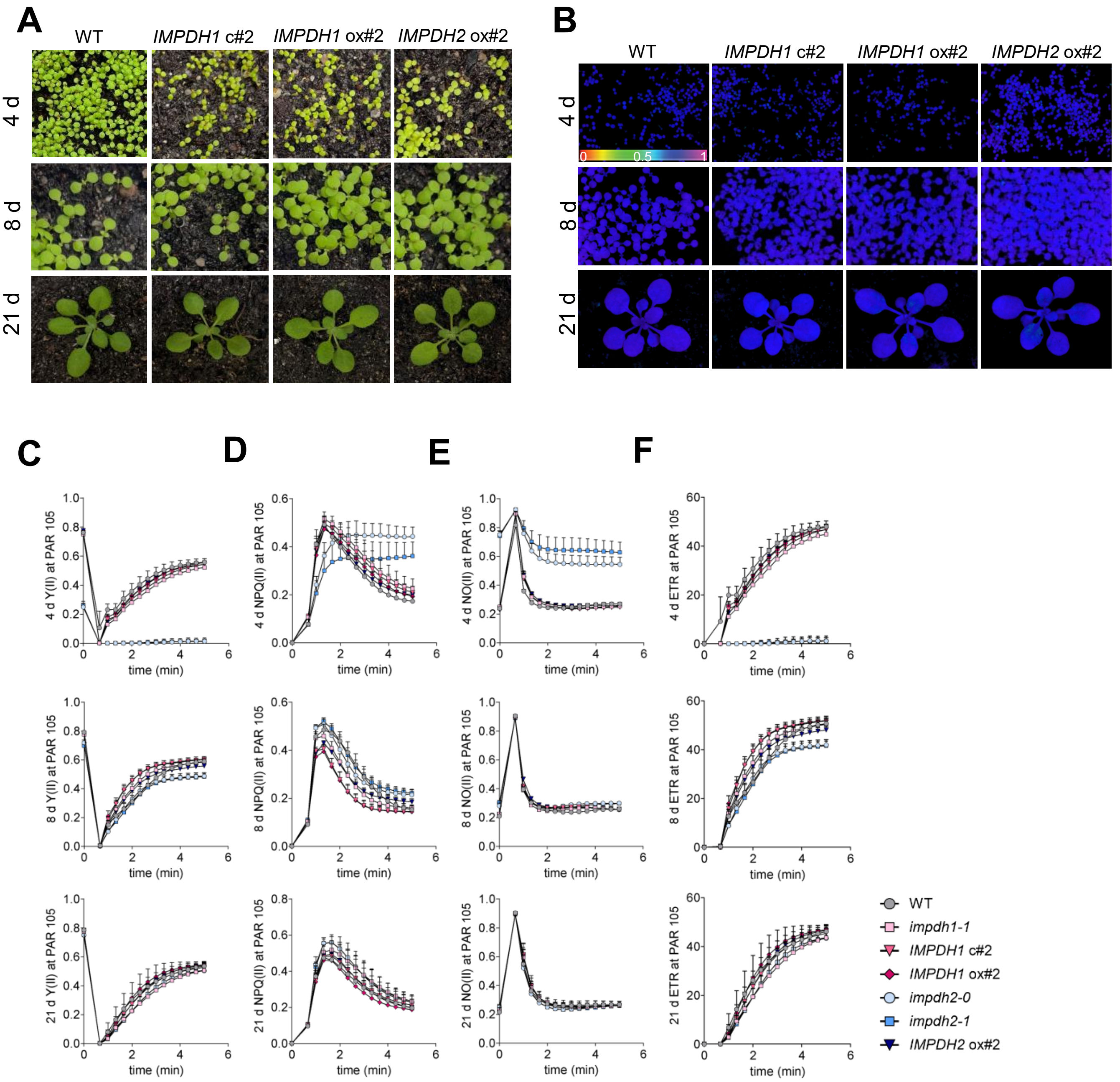


**Figure S7: Photosynthetic alterations in *IMPDH* mutants.** (**A**) Phenotypic analysis of WT, *IMPDH1* c#2, *IMPDH1* ox#2 and *IMPDH2 ox#2* Arabidopsis plants. Plants were grown under standard conditions (21 °C day and night temperature, 10 h day length and 120 *µ*E light intensity). (**B**) Representative PAM images of WT, *IMPDH1* c#2, *IMPDH1* ox#2 and *IMPDH2 ox#2* are depicted for PSII capacity (Fv/Fm). Light curves of Y(II) for WT, *impdh1*, *IMPDH1* c#2, *IMPDH1* ox#2, *impdh2-0*, *impdh2-1* and *IMPDH2 ox#2* after 4 d and 8d of growth. (**C-F)** Induction curves of Y(II), NPQ(II), NO(II), and ETR for WT, *impdh1*, *IMPDH1* c#2, *IMPDH1* ox#2, *impdh2-0*, *impdh2-1* and *IMPDH2 ox#2* after 4 d, 8d and 21d of growth.

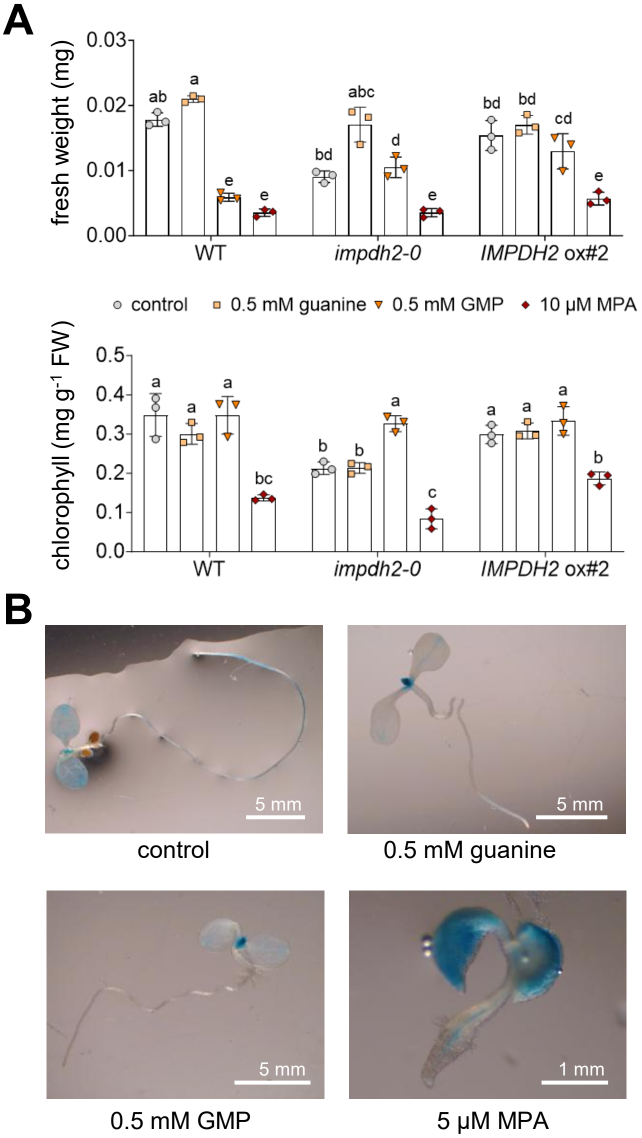


**Figure S8: Supplementation of IMPDH2 mutants.** (**A**) Fresh weight and chlorophyll contents of WT, *impdh2-0* and *IMPDH2* ox#2. Values are means ± SEM, n = 3 per treatment group. a-e, Means in a row without a common superscript letter differ ($P < 0.05$) as analyzed by two-way ANOVA and the TUKEY test. (B) Representative images of IMPDH2::GUS reporter line. The plants in (A-C) were grown in liquid MS without or with: 0.5 mM guanine, 0.5 mM GMP or 10 µM MPA for 6d grown under standard conditions (21 °C day and night temperature, 10 h day length and 120 *µ*E light intensity) on a shaker with 100 rpm.

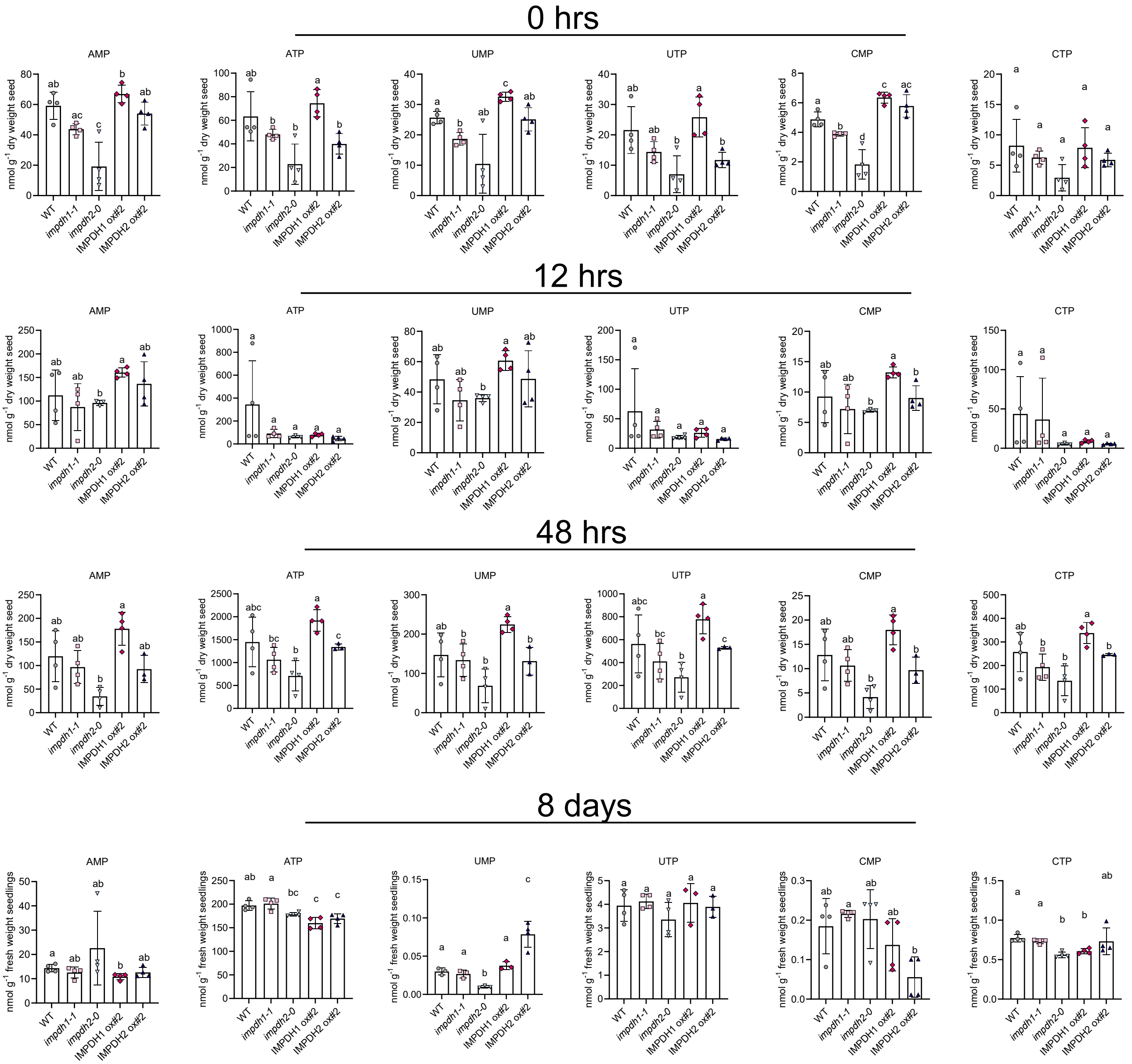


**Figure S9:** Metabolic analysis of guanylate biosynthesis pathway nucleotides in genetic IMPDH variants over a developmental time course. Nucleotides quantified by LC-MS after imbibition (0 h) and after 12 h, 48 h and 8 d of growth. Three to four biological replicates were analyzed, error bars are SD. The statistical analysis used the two-sided Tukey’s multiple pairwise comparison test with the sandwich variance estimator. Different letters indicate p values < 0.05.
